# Antiviral Sulfonated Polymersomes Inhibit Corona-, Influenza-, and Alphavirus

**DOI:** 10.64898/2026.10.11.758302

**Authors:** Sinead McCabe, Ruiqi Zhu, Raag Yagnik Setia, Jonathan C. Brown, Ziyi Pan, Xinyue Zhu, Xiaohao Xu, Beyzanur Kaya, Issabelle Wong, Fotoula Koropouli, Pedro S. Londoño, Kai Yi Mok, Ruthiran Kugathasan, Omar Rifaie-Graham, Vincenzo Torraca, Rui Pedro Galão, Wendy S. Barclay, Adrian Najer

**Affiliations:** Faculty of Life Sciences and Medicine, Institute of Pharmaceutical Science, King’s College London, London, UK; Faculty of Engineering, Department of Materials, Imperial College London, London, UK; Faculty of Medicine, Department of Infectious Disease, Imperial College London, London, UK; Faculty of Life Sciences and Medicine, Department of Infectious Diseases, King’s College London, London, UK; School of Physical and Chemical Sciences, Department of Chemistry, Queen Mary University of London, London, UK

**Keywords:** Polymersome, polymer nanoparticle, antiviral, COVID-19, flu, alphavirus

## Abstract

A broad-spectrum antiviral is required to enable rapid intervention against current, newly emerging and mutating viruses. Severe acute respiratory syndrome coronavirus 2 (SARS-CoV-2), influenza A virus (IAV), Sindbis virus (SINV) and chikungunya virus (CHIKV) all engage negatively charged moieties on host cells, such as heparan sulfate (HS) and sialic acid, to facilitate attachment for entry and infection. We designed HS-mimetic nanoparticles in the form of polymer vesicles (polymersomes) by blending membrane-forming and sulfonated copolymers. These biocompatible polymersomes successfully inhibited host cell infection of HS-dependent SARS-CoV-2 and HS-independent IAV in vitro with sub-nanomolar potency. Multivalent interaction of several virus ligands with a single polymersome as quantified by fluorescence correlation spectroscopy likely contributes to their potency. The sulfonated polymersomes also functioned in vivo as demonstrated by a reduction in the spread of SINV, an alphavirus model for CHIKV, in zebrafish embryos, which led to a significantly increased survival. Hence, our polymersomes could form the basis for an urgently required multifunctional broad-spectrum antiviral nanomedicine.

## Main

Viral pathogens create major concerns for human health, with viral emergence and mutations being a constant threat. Severe acute respiratory syndrome coronavirus 2 (SARS-CoV-2) is a prominent example of the threat viral infections pose. The COVID-19 pandemic resulted in over 7 million documented deaths globally and caused substantial socioeconomic disruption, including widespread economic losses and significant alterations to social structures worldwide.^1,2^ While the threat posed by SARS-CoV-2 has been substantially mitigated through widespread vaccination efforts, the persistent risk of viral mutation in SARS-CoV-2 and other emerging or yet unknown pathogens continues to present ongoing challenges for global health preparedness.^3,4^ Seasonal outbreaks of respiratory illness caused by influenza A virus (IAV) are another major contributor to global morbidity and mortality.^5^ Besides respiratory viruses, vector-borne viruses, such as chikungunya virus (CHIKV), have seen global resurgence in recent years.^6^ In 2025, over 400,000 cases of CHIKV were reported in 40 different countries.^7^

While some targeted antivirals exist for specific pathogens such as influenza viruses, HIV, and hepatitis viruses, no broad-spectrum antiviral therapeutics with demonstrated efficacy across diverse viral families are currently approved for clinical use, representing a critical gap in pandemic preparedness and treatment of emerging viral infections.^8^ Development of a broad-spectrum antiviral would enable pre-emptive protection against potential mutations as well as save valuable time and cost of developing a treatment once a pathogen has already posed a threat.^9^ The use of nanoparticles that directly interfere with viral entry represents one upcoming broad-spectrum antiviral strategy.^10^

Nanoparticle candidates for a broad-spectrum antiviral strategy predominantly aim to exploit conserved features of viral entry, e.g. electrostatic interactions of virions with anionic host cell surface moieties, such as heparan sulfate (HS) and/or sialic acid (SA).^11,12^ SARS-CoV-2, CHIKV, Sindbis virus (SINV), and many other viruses, utilize HS on the surface of host cells as the initial binding factor of the viral entry mechanism,^13^ while IAV represents a classical example that uses SA binding instead of HS.^14^ Hence, a range of nanomaterial architectures that present or mimic HS/SA have been developed as broad-spectrum antivirals, including inorganic nanoparticles,^15–17^ polymer nanoparticles,^18–20^, lipid nanoparticles (LNPs),^21–23^ dendritic nanoparticles,^24–26^ and nanosheets.^27,28^ Interestingly, sulfated/sulfonated nanoparticles that mimic HS were previously found to extend their inhibitory potential also to HS-independent, SA-binding viruses, such as IAV, revealing an even broader target range beyond HS binders.^29^

Despite the broad-spectrum potential of HS/SA nanoparticles against various viruses, development towards clinical translation remains limited by very few in vivo efficacy studies. Depending on the construct there are either concerns about low potency, reversible binding mechanism, or biocompatibility, while routes of administration and susceptibility to viral mutation are other important factors that require attention.^10–12^ Possible approaches to overcome some of the challenges associated with antiviral nanoparticles include the design of virucidal nanoparticles,^16^ co-delivery of antiviral compounds,^30^ or coordination with the immune system.^31,32^ Polymer vesicles made from block copolymers, called polymersomes,^33^ represent a promising nanoparticle platform for these three approaches given their membranous nature and availability of a hydrophilic core and a hydrophobic membrane for varied cargo loading. Analogous vesicles made from lipids (liposomes) have recently allowed formulation of virucidal systems that function via membrane fusion,^21,22^ while polymersomes have previously been designed for co-delivery of antiviral compounds.^30,34^ Therefore, developing a potent, broad-spectrum, virion-binding nanoparticle based on the very versatile polymersome platform will unlock alternative possibilities for virucidal systems, co-delivery of antiviral drugs, and immune cell coordination.

We herein developed and tested copolymer-based HS-mimicking nanoparticles in the form of micelles and polymersomes to competitively bind to viral surface proteins to inhibit interaction between virus and host cells, thereby preventing infection. We first utilized our previously identified chemical modification strategy on a poly(acrylic acid) (PAA) containing copolymer^18,35^ to enable sulfonation of other copolymers. We then co-assembled these copolymers with vesicle-forming, inert poly(ethylene glycol) (PEG) containing copolymers to create antiviral HS-mimicking polymersomes. The obtained HS-mimetic nanoparticles were first tested against a SARS-CoV-2 pseudovirus (PV) system, which revealed higher potency of the polymersomes versus micelles. The high potency of polymersomes can be related to multivalent interaction with SARS-CoV-2 spike proteins as revealed by fluorescence correlation spectroscopy (FCS). The clinical potential and broad-spectrum applicability of sulfonated polymersomes were demonstrated by inhibiting whole infectious SARS-CoV-2 and HS-independent IAV in vitro. Final in vivo evaluation of the polymersomes was performed with a Sindbis virus (SINV) zebrafish embryo infection model, which represents an alphavirus model for CHIKV. The polymersomes reduced the spread of the virus and increased survival of the embryos, revealing their potential for further development towards an urgently required broad-spectrum antiviral treatment.

Polymeric nanoparticles have been chosen as the nanoparticle platform to create HS-mimetic viral inhibitors due to the potent virus-binding capabilities of polymers. Self-assembled polymersomes were expected to create efficient virus inhibitors given their high flexibility and size commensurate with virions (70-120 nm). These are two previously identified characteristics that can improve virus entry inhibition potency using other nanomaterial classes.^25,36^ Polymer-based HS-mimetic micelles were first formulated using amphiphilic block copolymers poly(butadiene)-*block*-poly(acrylic acid) (PBD-PAA), poly(butyl acrylate)-*block*-poly(acrylic acid) (PBA-PAA), and poly(dimethylsiloxane)-*block*-poly(acrylic acid) (PDMS-PAA), analogously to our previously published poly(_DL_-lactic acid)-*block*-poly(acrylic acid) (PDLLA-PAA) micelles.^18^ The hydrophilic PAA on each copolymer has repetitive carboxylic acid units that were chemically modified with a small sulfonate containing molecule, 5-amino-2-methoxybenzenesulfonic acid (AMBS),^18,35^ to create the HS mimicking properties of the nanoparticles (Figure 1a). AMBS modification was carried out in suspension after micellar self-assembly using direct dispersion or the solvent injection method, as published previously for our PDLLA-PAA-based micelle system.^18^ Sulfonation of micelles (PBD-M, PBA-M, PDLLA-M, and PDMS-M) was confirmed using the Farndale assay (Figure S4), which enabled quantification of polysulfonate concentration in the final micelle preparations,^37,38^ which was also used to calculate antiviral potency subsequently. HS-micelles were characterized through dynamic light scattering (DLS) and zeta potential measurements. PBD-PAA-AMBS and PBA-PAA-AMBS (PBD-M and PBA-M) revealed hydrodynamic diameters of 15.4 ± 0.9 nm and 26.0 ± 2.1 nm, respectively (Figure 1b-e). The zeta potentials were anionic as expected, which was -28.8 ± 0.6 mV for PBD-M and -33.6 ± 1.8 mV PBA-M (Figure 1g,h). PDLLA-PAA and PDMS-PAA micelles had similar zeta potential values (Figure S1d) compared to the PBD-/PBA-based micelles.

**Figure 1.**
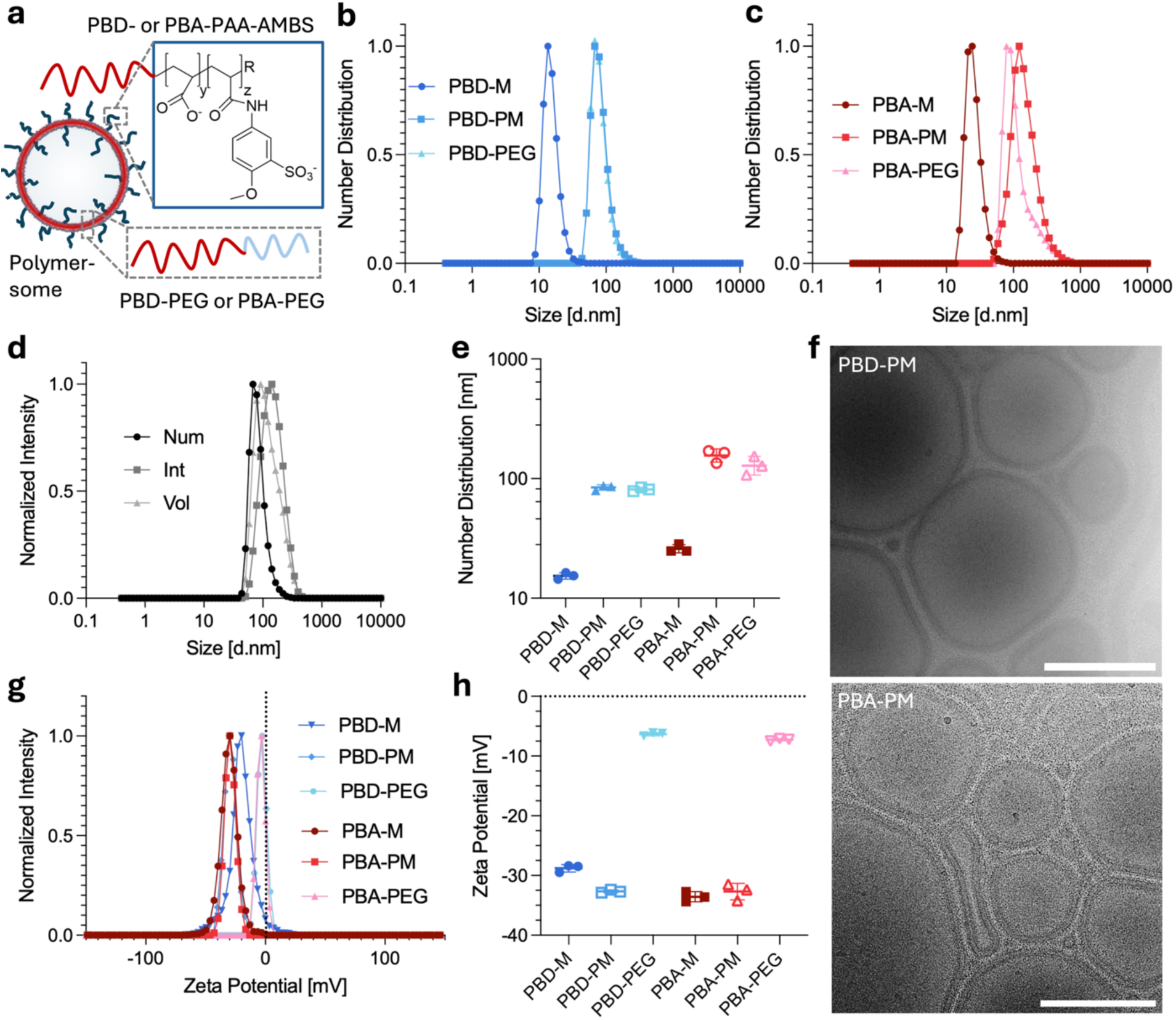
HS-micelles and polymersomes characterization. (a) Polymersome schematic showing the blends of PBD-PEG/PBD-PAA-AMBS and PBA-PEG/PBA-PAA-AMBS that were developed. (b,c) Representative DLS curves (number distributions) for PBD-PAA-AMBS/PBA-PAA-AMBS micelles and corresponding polymersomes made from PBD-PEG or PBA-PEG with and without 10 mol% PBD-PAA-AMBS or PBA-PAA-AMBS (PBD-PM/PBA-PM), respectively (average of technical triplicates). (d) Average DLS size distribution (intensity, volume, and number distribution) of sulfonated polymersome PBD-PM (technical triplicates, curves for PBA-PM can be found in Figure S1a). (e) Average DLS number distribution (mean ± SEM, technical triplicates). (f) Cryo-TEM images of PBD-PM and PBA-PM. Scale bars are 200 nm. (g,h) Representative zeta potential curves (average of technical triplicates) and average values (mean ± SEM, technical triplicates) for HS-micelles, HS-polymersomes, and respective control polymersomes.

To formulate polymersomes (PM) with HS-mimetic characteristics, the modified block copolymers of interest, PBD-PAA-AMBS and PBA-PAA-AMBS, were ion-exchanged and freeze-dried to allow mixing at different ratios in organic solvent (ethanol) with either poly(butadiene)-*block*-poly(ethylene glycol) (PBD-PEG) or poly(butyl acrylate)-*block*-poly(ethylene glycol) (PBA-PEG), respectively, to assemble PBD-PM and PBA-PM using the film rehydration technique. PBD-PEG and PBA-PEG are two readily employed copolymers for polymersome formation,^39–41^ while polymersomes made from blends of PBD-PEG with unmodified PBD-PAA were previously achieved for applications beyond antivirals.^42^ The addition of mostly PEGylated vesicle-forming copolymers was hypothesized to promote co-assembly into bilayers incorporating a smaller fraction of anionic block copolymer. In turn, high content of the micelle-forming PBD-PAA-AMBS and PBA-PAA-AMBS in the PM formulations was expected to disrupt vesicle formation and yield micelles instead. PM formulations were initially validated with the use of highly hydrophilic sulforhodamine B (SRB) dye encapsulation experiments. SRB encapsulation should only be possible with PM morphology because micelles lack an aqueous core for entrapment. The goal was to maximize the number of PBD-PAA-AMBS and PBA-PAA-AMBS chains within our PM formulations to increase the chance of multivalent interactions with virions. 10 mol% of the modified, anionic copolymers were identified as a good compromise of high ligand content while still mainly forming polymersomes (Figure S2a). This agrees well with a previous study on blending PBD-PEG and unmodified PBD-PAA, where 10 mol% was also found to be the border for formulating structurally intact polymersomes.^42^ Hence, 10 mol% of active copolymer was kept constant for all subsequent polymersome experiments for PBD-PM and PBA-PM.

Sulfonation of the final HS-polymersomes was again quantified with the Farndale assay. HS-polymersomes were further characterized by DLS revealing a diameter of around 100 nm for PBD-PM and 150 nm for PBA-PM (Figure 1e and Figure S1a,b). The zeta potential for HS-polymersomes were dominated by the anionic copolymer resulting in lower zeta potentials compared to the PEG-only polymersome controls (Figure 1g,h). The absence of a second neutral peak for the HS-polymersomes is an indicator that the copolymers mixed well. Cryogenic transmission electron microscopy (cryo-TEM) confirmed the membranous morphology of both PBD-PM and PBA-PM at 10 mol% anionic copolymer integration (Figure 1f and Figure S2b) as expected from the SRB encapsulation experiments (Figure S2a). In conclusion, a set of different polymeric nanoparticles has been developed successfully, consisting of sulfonated micellar controls, sulfonated polymersomes that incorporate 10 mol% of the modified micelle-forming copolymer, and PEGylated control polymersomes, which enabled detailed analysis of their antiviral potential.

The antiviral mechanism of sulfonated polymersomes is expected to be based on multivalent interaction of multiple viral ligands with the polymersomes as is the case for most antiviral HS/SA mimetic nanoparticles.^11^ The interaction of SARS-CoV-2 spike protein with HS-mimetic copolymers embedded within our polymersome membranes was expected given the known binding affinity of HS/heparin with this target.^43,44^ We employed single-molecule sensitive fluorescence correlation spectroscopy (FCS) to characterize the binding interactions between HS-polymersomes and fluorescently labelled SARS-CoV-2 spike protein (Spike-488, Figure 2a). FCS has previously been used extensively to study protein interactions with polymersome surfaces,^39,45,46^ including multivalent binding of pathogen ligands.^47^ Autocorrelation analysis revealed distinct Spike-488 binding behaviors between the different formulations tested (Figure 2b,c). Both HS-polymersome formulations demonstrated significantly altered diffusion properties compared to Spike-488 alone, indicating partial formation of larger bound complexes upon mixing. The percentage of particles exhibiting slow diffusion characteristics was only increased for PBD-PM and PBA-PM formulations compared to the PEG-only controls (PBD-PEG and PBA-PEG) as obtained from the two-component fits (Figure 2d). Control experiments with fluorescently labelled human serum albumin (HSA) did not show an increase in the particle fraction for any of the samples (Figure S3), confirming that the interaction of Spike-488 with PBD-PM and PBA-PM is specific.

**Figure 2.**
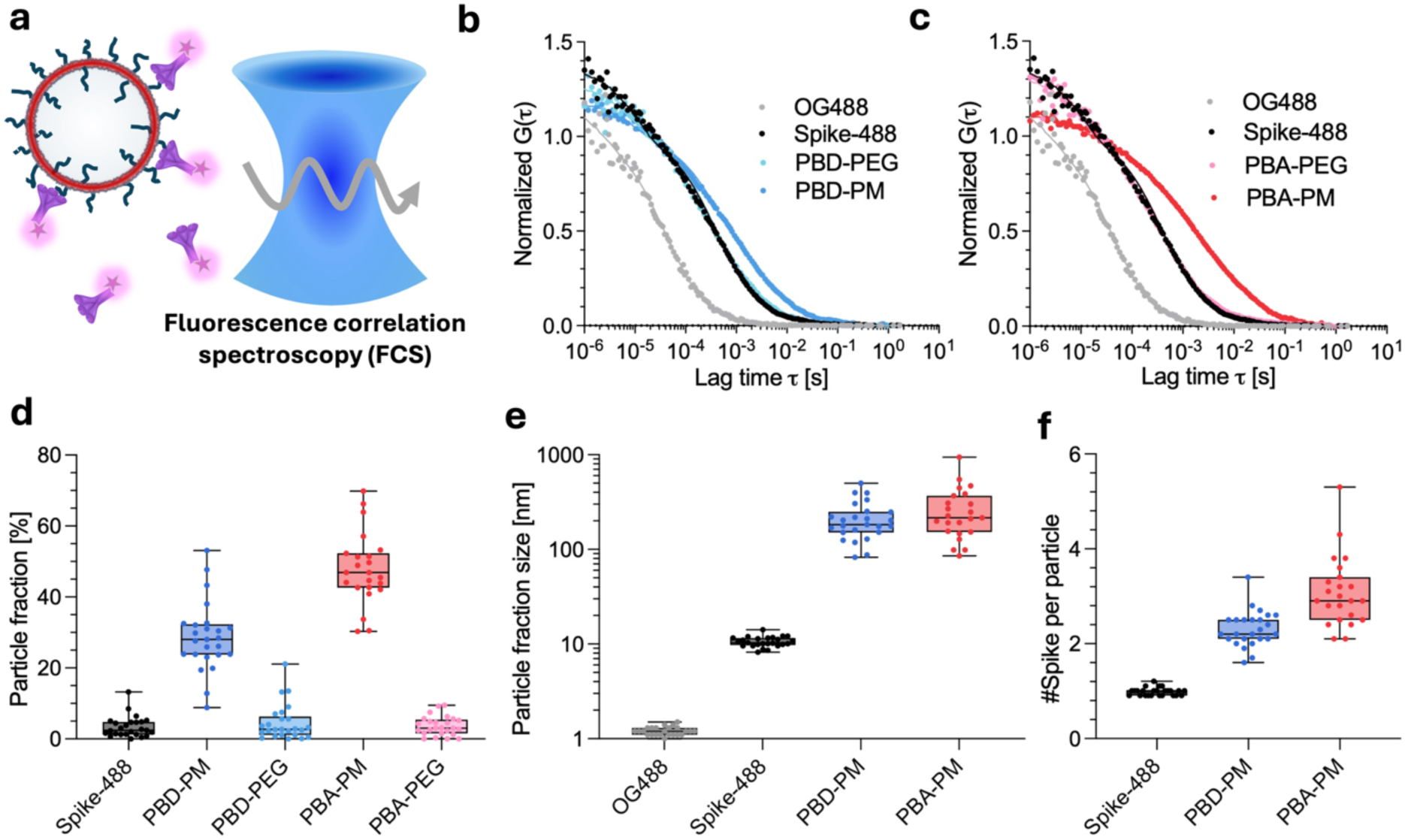
Fluorescence correlation spectroscopy analysis of HS-polymersome-spike protein interaction. (a) Schematic representation of FCS experimental setup. Schematic modified from Servier Medical Art website CC-BY. (b,c) Average normalized autocorrelation curves for SARS-CoV-2 AF488-labeled spike protein (Spike-488, BA.2 Spike GCN4-IZ) and corresponding mixtures of Spike-488 with PBD-PEG, PBA-PEG, PBD-PM and PBA-PM (*n* = 25 measurements). (d,e) Quantification of particle fraction and hydrodynamic diameter for the slow diffusing species obtained from two-component fits of data in b,c (*n* = 25 measurements). Reference values for OG488 and Spike-488 data were obtained from one component fits. (f) Calculation of number of Spike-488 bound per slow diffusing particle using molecular brightness data (counts per particle, cpp) from the autocorrelation analysis in b,c (*n* = 25 measurements). Box-plots: Centre line, the median; box limits, upper and lower quartiles; whiskers, minimum and maximum values.

Analysis of FCS-derived hydrodynamic particle size revealed substantial increases for Spike-488 upon HS-polymersome binding (Figure 2e). Free spike protein exhibited a hydrodynamic diameter of 10.6 ± 1.3 nm, which increased about 20-fold for the particle fractions of the mixtures of Spike-488 with PBD-PM and PBA-PM. The obtained sizes correspond well to the sizes obtained from DLS intensity distributions of the particles only (Figure 1d and Figure S1a). The number of spike proteins per particle was calculated by comparing molecular brightness (counts per particle, cpp) of the particle fractions from the mixtures versus cpp of Spike-488 alone (Figure 2f). PBD-PM and PBA-PM could both bind multiple Spike-488. These data indicate successful interaction of multiple spike proteins with individual HS-polymersomes, forming multimeric complexes, supporting their potential as multivalent antiviral virus-binding agents. The FCS data collectively demonstrate that the functional, anionic copolymer is essential for polymersomes to successfully bind SARS-CoV-2 spike protein.

After demonstrating successful multivalent SARS-CoV-2 spike protein binding to HS-polymersomes, we first selected a pseudovirus (PV) in vitro model for antiviral testing as it is replication deficient, enabling specific assessment of inhibitor action on viral entry.^48^ We utilized a SARS-CoV-2 PV system with a luciferase gene for simple luminescence readout and expression of the D614G spike protein,^49^ to test the entry inhibition potential for our HS-micelles and HS-polymersomes (Figure 3a). We first compared efficacy among sulfonated micelles prepared with three different copolymers, in comparison with our previous PDLLA-PAA-AMBS system to evaluate whether the inhibition potential could be transferred to PBD-, PBA-, and PDMS-based copolymers with the same PAA hydrophilic block (Figure 3b,c).^18^

**Figure 3.**
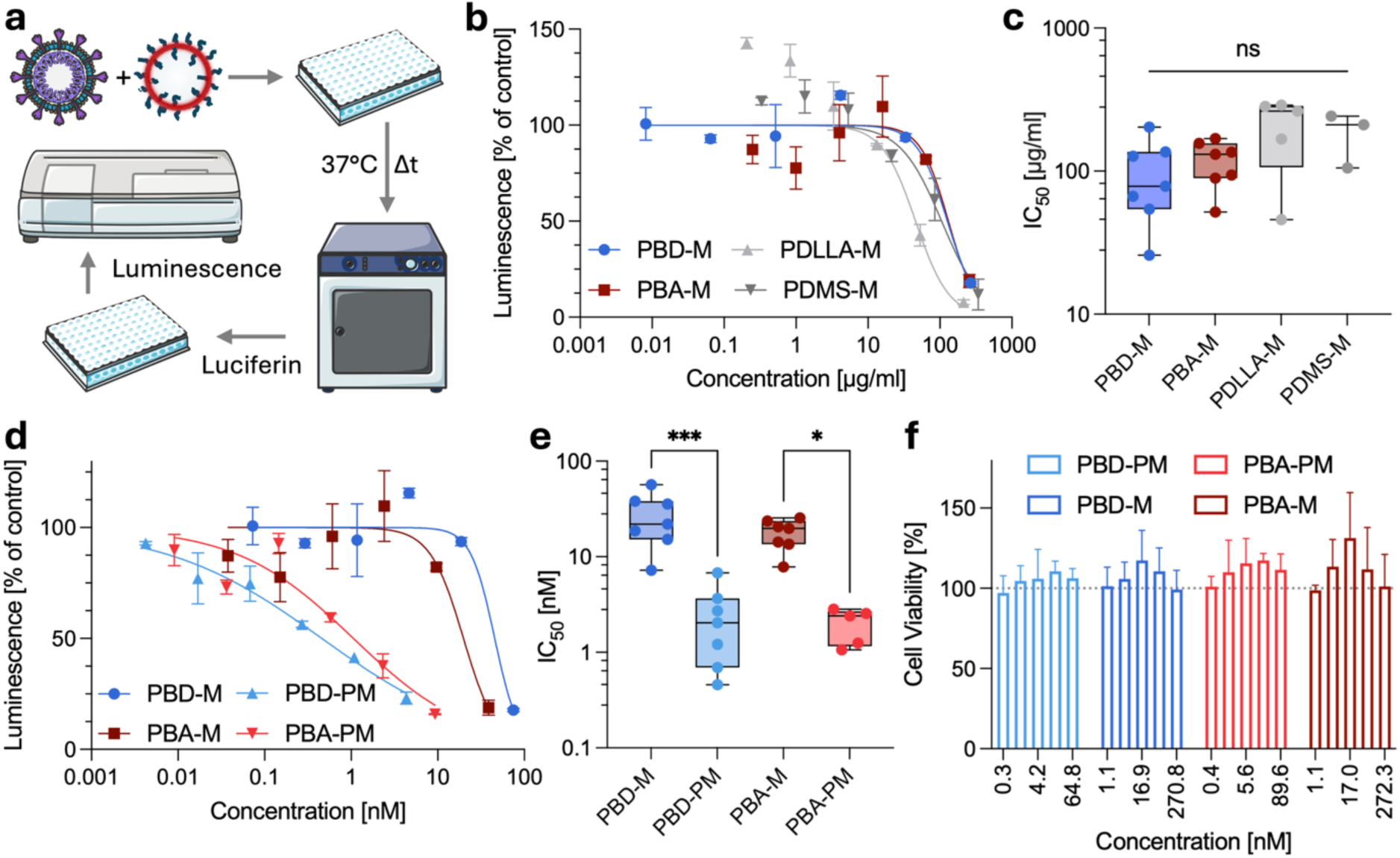
HS-mimicking polymeric micelles and polymersomes inhibit SARS-CoV-2 PV entry into HeLa-ACE2 cells. (a) Schematic of the PV inhibition assay workflow. HS-micelles or polymersomes were pre-incubated with PV before addition to HeLa-ACE2 cells, followed by 48 h incubation and luminescence readout. Schematic modified from Servier Medical Art website CC-BY. (b) Representative dose–response curves showing luminescence as a function of functional polymer concentration from Farndale assays (µg/ml) for sulfonated PBD-M, PBA-M, PDLLA-M, and PDMS-M (mean ± range for *n* = 1 independent experiment with technical duplicates). (c) IC₅₀ values (µg/ml) for each micelle formulation derived from dose-response curves as shown in b (*n* ≥ 3 independent repeats with technical duplicates). One-way ANOVA with Tukey’s multiple comparison test. ns = not significant. (d) Dose-response curves for PBD-and PBA-based micelles versus polymersomes against SARS-CoV-2 D614G PV in HeLa-ACE2 cells (mean ± range for *n* = 1 independent experiment with technical duplicates). (e) IC₅₀ values (nM, nanoparticle concentration) derived from PV dose-response fits for micelles (data converted to nM from c) as shown in d (*n* ≥ 3 independent repeats with technical duplicates). One-way ANOVA with Sidak’s multiple comparison test, *=p<0.05; ****=p<0.0001. (f) Cell viability of HeLa-ACE2 cells incubated with HS-mimetic nanoparticles for 48 h (*n* ≥ 3 independent repeats with technical duplicates). Box-plots: Centre line, the median; box limits, upper and lower quartiles; whiskers, minimum and maximum values.

All four HS-micelle formulations achieved a reduction in the luminescence signal in a concentration-dependent manner confirming PV entry inhibition potential for all (Figure 3b). IC₅₀ values derived from dose-response curves revealed similar mean values of about 100 to 200 µg/mL, with no significant differences among the four different copolymers (Figure 3c). Hence, keeping the functional PAA-AMBS block constant while varying the backbone hydrophobic polymers that form the micellar core from PDLLA to PBD, PBA, or PDMS produced nanoparticles with similar viral entry inhibition efficacy. Hence, the AMBS modification procedure and inhibition potential could successfully be transferred from our previously published PDLLA-M^18^ to other micelle-forming copolymer building blocks. All the HS-micelles were tested for cytocompatibility with the same host cells as used in the above PV assays (HeLa-ACE2) and at the same concentrations for the same time frame (Figure 3f and Figure S5b). None of the micelles showed any cell toxicity under these conditions, highlighting their potential for further development towards more potent nanoparticle formulations that contain these copolymers as one component.

Polymersomes that contain 10 mol% of PBD-PAA-AMBS or PBA-PAA-AMBS were next tested in the same PV in vitro model (Figure 3d,e). To allow direct comparison of inhibition potential on a particle concentration basis, FCS was employed to quantify nanoparticle concentrations of micelles and polymersomes by labelling with a hydrophobic fluorescent dye at a fixed functional polymer concentration obtained from Farndale assays.^18^ This then allowed back and forth conversion of nanoparticle concentration (nM) and functional polymer concentration (µg/mL). In the PV assay, both sulfonated polymersome formulations inhibited infection in a concentration-dependent manner at low nanomolar concentrations (Figure 3d,e). The micellar versions, PBD-M and PBA-M, showed mean IC₅₀ values (Figure 3c), calculated from dose response curves (Figure 3b) after conversion to particles numbers (Figure 3e), of 27.7 ± 6.4 nM (equals 99 ± 23 µg/mL) and 17.9 ± 2.4 nM (equals 118 ± 16 µg/mL) , respectively. The HS-polymersome formulations were markedly more potent with PBD-PM achieving a mean IC₅₀ of 2.5 ± 0.8 nM (equals 10.3 ± 3.4 µg/mL functional polymer concentration), representing around a 16-fold improvement over PBD-M, while PBA-PM reached 2.0 ± 0.4 nM (equals 20.8 ± 3.7 µg/mL), around an 8-fold improvement over PBA-M. Importantly, also the HS-polymersomes were cytocompatible to the HeLa-ACE2 cells (Figure 3f), confirming that the observed inhibition reflects a specific viral entry inhibition effect rather than non-specific cytotoxicity. The higher inhibition potential of polymersomes versus micelles can be explained by their difference in size. The micelles are substantially smaller (ca. 15 – 25 nm, Figure 1b-e) than the virions (80 – 120 nm) they are intended to inhibit. In agreement with our results, maximal inhibition efficacy of virus-size matching nanoparticles was identified before using inorganic nanoparticles.^36^ Flexibility of the inhibitor was previously found as another important factor determining efficacy, with more flexible structures showing higher inhibition efficacy.^25,50^ Polymersomes made from low glass transition temperature (*T_g_*) copolymers, such as PBD-and PBA-based copolymers, are known to exhibit high flexibility and fluidity given their dynamic membrane architecture with an aqueous core,^51^ which could be another contributor to their high efficacy of viral inhibition. The most potent inhibitors identified up to now, the HS-polymersomes, were taken forward for in depth testing of their antiviral potency in vitro and in vivo.

HS-polymersomes were further tested against whole infectious SARS-CoV-2 (Figure 4a-c) to confirm their inhibition potential found with the PV model (Figure 3). The experiment was performed analogously to the PV trials but now using Vero E6 host cells and the Omicron variant of SARS-CoV-2 (BA.1 lineage, isolate M21021166). After 48 hours of incubation, intracellular SARS-CoV-2 nucleocapsid protein expression was quantified by ELISA as a measure of infection. Both HS-polymersome formulations exhibited potent inhibition of whole infectious SARS-CoV-2 (Figure 4b-c) as expected from the PV data. The PBD-PM had a IC_50_ value of 0.38 ± 0.15 nM (1.6 ± 0.6 µg/mL) and the PBA-PM had a IC_50_ value of 0.21 ± 0.04 nM (2.2 ± 0.4 µg/mL) without any cytotoxicity observed in these host cells either (Figure S5a). Control polymersomes without the sulfonated copolymer (PBD-PEG and PBA-PEG) did not inhibit viral infection (Figure S5c). The potent inhibition potential of PBD-PM and PBA-PM can be explained by the strong HS binding characteristic of Omicron due to positively charged amino acids in its spike protein^23^ making this variant highly susceptible to inhibition with anionic antiviral constructs.^52–54^ Polymeric nanoparticles, liposomes and LNPs are most closely related to our polymersome design given their similar size and flexible nature. Recent tests with antiviral LNPs that contain sulfonated components (POSTAN) showed IC_50_ values at 101 µg/mL of their active component against Omicron.^23^ Sulfonated liposomes were tested against Omicron PV, with IC_50_ of 51 µg/mL.^22^ Hence, our sulfonated HS-polymersomes at 1.6 – 2.2 µg/mL IC_50_ were at least one order of magnitude more potent inhibitors of SARS-CoV-2 Omicron compared to these similar nanosystems with the caveat of assay differences. Broad-spectrum applicability was next demonstrated by inhibiting IAV (A/England/195/2009(H1N1)).

**Figure 4.**
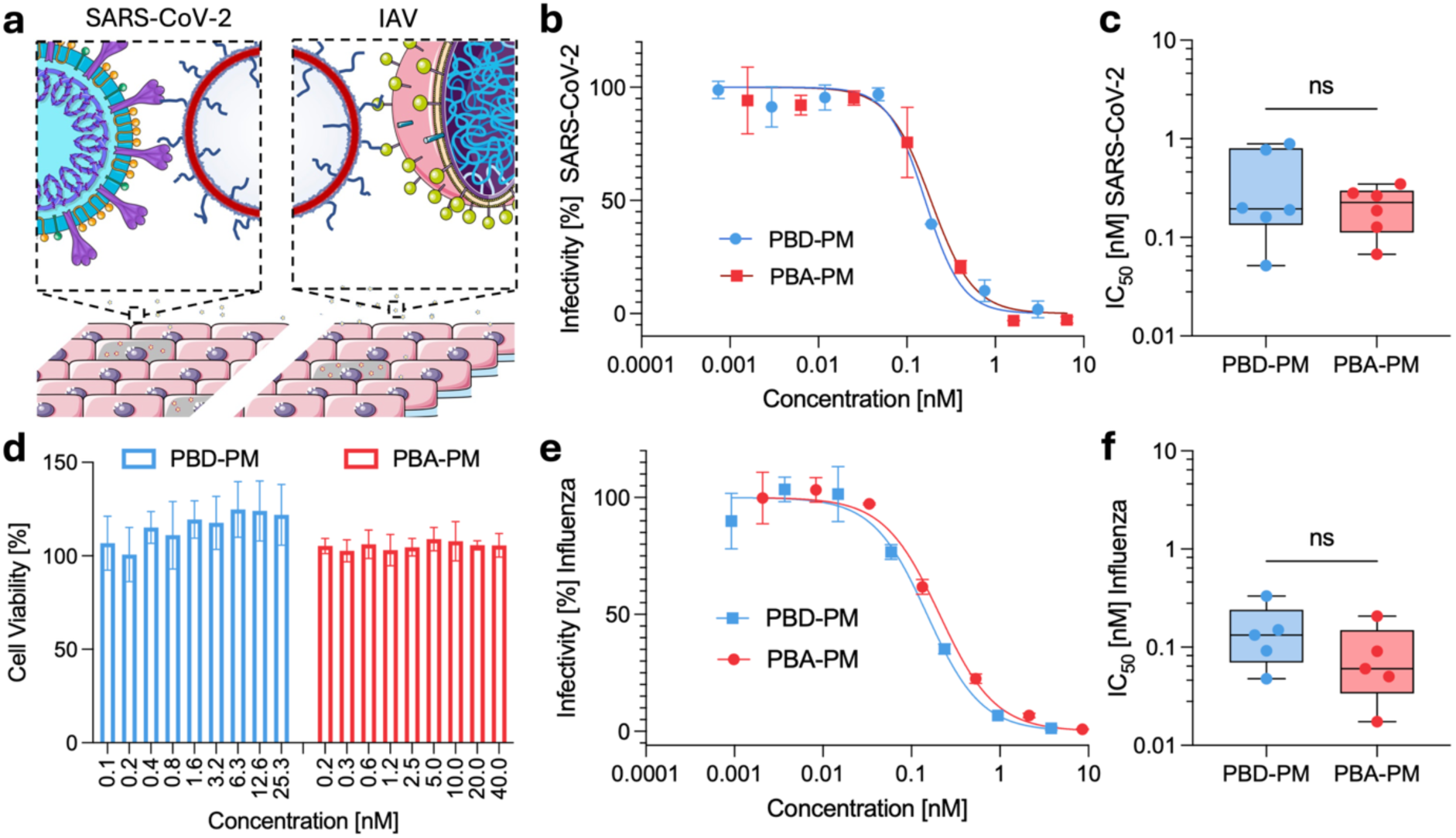
HS-polymersomes show in vitro activity against whole infectious SARS-CoV-2 and IAV. (a) Schematic depicting the proposed mechanism of action. HS-mimicking polymersomes act as decoys, competitively binding to the virions and blocking engagement with cell-surface receptors, thereby preventing viral entry. Schematic modified from Servier Medical Art website CC-BY. (b) Representative dose–response curves showing infectivity of SARS-CoV-2 Omicron BA.1 lineage (isolate M21021166) as a function of nanoparticle concentration (nM) for sulfonated PBD-PM and PBA-PM (mean ± range for *n* = 1 independent experiment with technical duplicates). (c) IC₅₀ values (nM, nanoparticle concentration) derived from whole infectious virus dose-response fits for polymersomes as shown in b (*n* ≥ 3 independent repeats with technical duplicates). Two-tailed Mann–Whitney test, ns = not significant. (d) Cell viability of MDCK cells treated with increasing concentrations of PBA-PM and PBD-PM, measured by MTS/PMS assay and expressed relative to untreated controls after 18 hr (*n* ≥ 3 independent repeats with technical duplicates). (e) Representative dose–response curves showing IAV (A/England/195/2009(H1N1)) infectivity in MDCK cells by PBD-PM and PBA-PM. (f) IC₅₀ values for PBD-PM and PBA-PM inhibition of IAV derived from dose-response curves as shown in e. Unpaired t-test, ns = not significant. Control data for PBD-PEG and PBA-PEG can be found in Figure S5c. Box-plots: Centre line, the median; box limits, upper and lower quartiles; whiskers, minimum and maximum values.

IAV does not engage HS but instead binds to another anionic moiety, SA, for cell entry and infection.^14^ The anionic nature of both HS and SA explains why previous HS-mimetic nanoparticles were found to be active against IAV despite not intentionally designing the inhibitors for this application.^29^ Motivated by these previous results, we wanted to test whether our PBD-PM and PBA-PM also have the ability to neutralize whole infectious IAV (H1N1) and prevent infection of Madin-Darby Canine Kidney (MDCK) cells. Both HS-polymersomes produced a concentration-dependent reduction in viral infectivity (Figure 4e). This was reflected in the derived IC₅₀ values of PBD-PM at 0.15 ± 0.05 nM (0.62 ± 0.20 µg/mL) and 0.085 ± 0.033 nM (0.88 ± 0.34 µg/mL) for PBA-PM (Figure 4f). To confirm that the loss of infectivity reflected viral inhibition rather than cytotoxic effects on the host cells, cytocompatibility assays were performed in MDCK cells across a concentration range that extended well above the measured IC₅₀ (Figure 4d). Neither PBD-PM nor PBA-PM reduced viability with cells remaining above 80% viability at every concentration tested. The absence of toxicity at concentrations far exceeding the IC₅₀ establishes a wide therapeutic window and confirms that the reduction in infectivity is attributable to inhibition of viral cell entry rather than to compromised cell health. A leading sulfonated nanoparticle in the literature (MUS:OT-NP) has previously shown EC_50_ at 1.4 µg/mL equaling 6.3 nM against H1N1.^29^ Again, our polymersomes are more potent in comparison. Interestingly, polystyrene sulfonate (PSS), which is structurally related to our functional polymers did not inhibit IAV.^17^ Our PAA-AMBS on the surface of our polymersomes presents aromatic sulfonates similar to PSS. However, due to the high density of functional groups in PAA we only modify about 30% of the carboxylic acid groups with our synthesis protocol,^18^ creating a mixture of aromatic sulfonates and carboxylic acids in the final product, which could explain the difference in IAV inhibition compared to PSS.

To continue to assess the antiviral efficacy of our HS-polymersome formulations in a physiologically relevant in vivo model, we selected a zebrafish embryo model of SINV infection.^55^ SINV is an alphavirus that recapitulates key features of CHIKV pathogenesis in zebrafish embryos, providing an ethically accessible vertebrate model for evaluating antiviral interventions.^56,57^ Zebrafish embryo infection models have been established as versatile platforms for studying a range of viral infections^58^ and for evaluating the in vivo efficacy of antiviral candidates.^59^ Both SINV and CHIKV are HS-dependent and can be inhibited with HS/heparin.^59,60^ The POSTAN LNPs previously showed similar inhibition of SARS-CoV-2 Omicron and CHIKV, which served as a rational to test our polymersomes in a related SINV in vivo model.^23^ Zebrafish embryos were infected with SINV (20 PFU) pre-incubated with either PBS or HS-polymersomes for one hour prior to injection into the vasculature via the dorsal aorta at 2 days post-fertilization. Survival and infection severity were monitored until 72 hours post-infection (hpi) (Figure 5).

**Figure 5.**
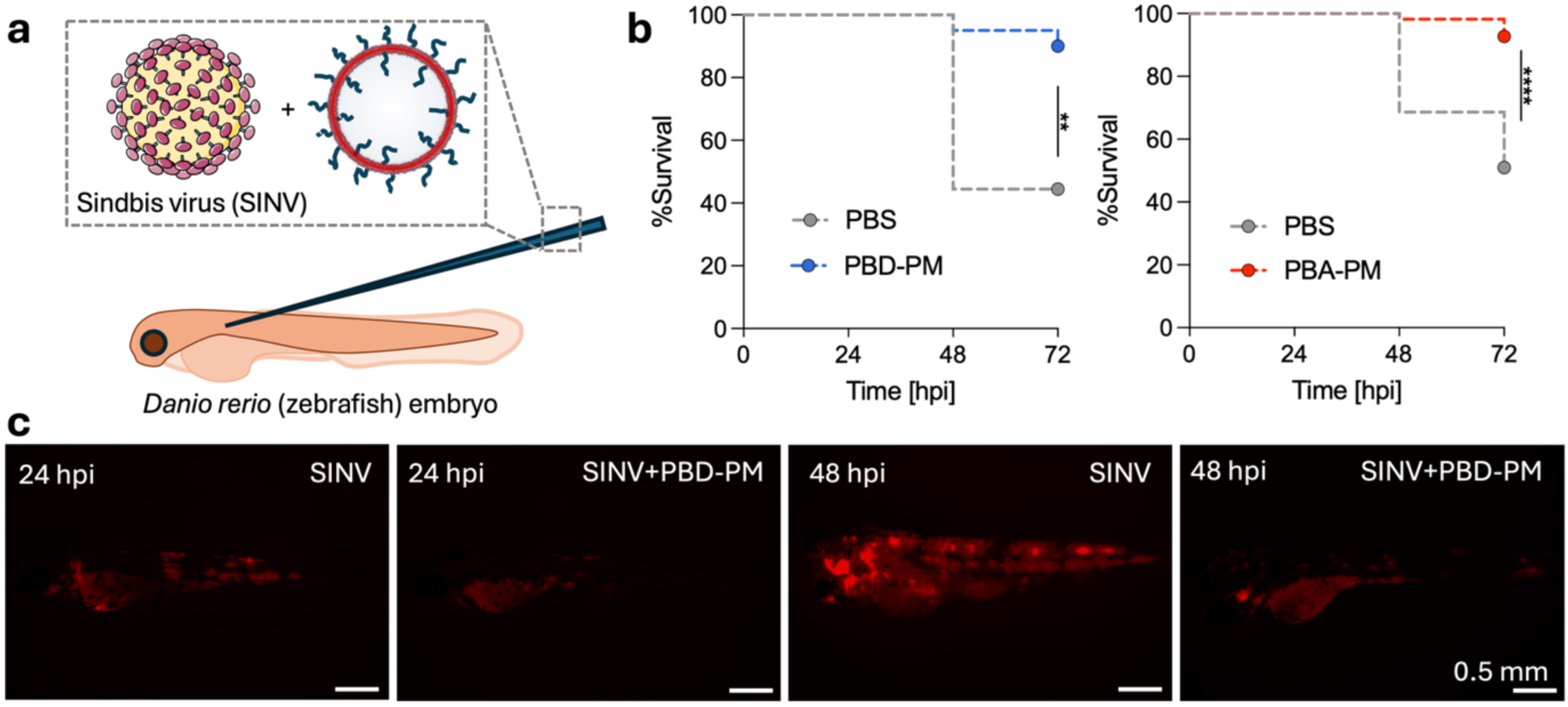
Sindbis virus (SINV) inhibition by polymersomes in zebrafish embryo model. (a) Schematic of in vivo experiment modified from Servier Medical Art website CC-BY. (b) Kaplan–Meier survival curves of zebrafish embryos following injection with SINV (20 PFU) pre-incubated with PBS or PBA-PM or PBD-PM. Survival was monitored at 24, 48, and 72 hpi (*n* = 1 or 3 independent cohorts of ∼20 larvae per condition for PBD-PM and PBA-PM, respectively). Curve comparison with Log-rank (Mantel-Cox) test, **p<0.01; ****p<0.0001. (c) Representative fluorescence micrographs of zebrafish embryos at 24h hpi and 48 hpi showing SINV dissemination (red) following injection with SINV + PBS or SINV + PBD-PM. Scale bars, 0.5 mm.

Survival analysis of the zebrafish experiments confirmed a significant protective effect for both HS-polymersome formulations (Figure 5b,c). In PBS-infected controls, survival dropped to approximately 45% and 50% by 72 hpi. By contrast, embryos receiving SINV pre-incubated with and PBD-PM-treated embryos similarly sustained ∼90% survival at 72 hpi and PBA-PM maintained ∼95% survival over the full 72-hour observation period. Direct visualization of viral dissemination by fluorescence microscopy further supported the protective effect of the HS-polymersomes (Figure 5c). At 24 and 48 hpi, SINV infection in PBS-control embryos had a clear detectable fluorescent signal, which had spread throughout the larval body. In contrast, PBD-PM treated embryos displayed markedly restricted viral replication at the 48 hpi timepoint, with infection remaining substantially more localized and of lower intensity compared to PBS controls. Together, these data demonstrate that PBD-PM and PBA-PM can suppress viral infectivity in vivo, reducing disease severity by restricting viral replication and dissemination, conferring significant protection from virus-induced lethality in the zebrafish embryo model. The retention of inhibitory activity in an in vivo context is promising, as it suggests that PBA-PM and PBD-PM are sufficiently stable and active under physiological conditions to confer meaningful protection at the whole-organism level using this prophylactic scenario. Taken together, these findings establish PBD-PM and PBA-PM as lead formulations amongst our candidates, combining sub-nanomolar antiviral potency against whole infectious virus, comparable activity across HS-dependent and HS-independent viruses, and in vivo efficacy.

In conclusion, we have shown that sulfonated micelles with different backbone hydrophobic polymers produce similar antiviral candidates. By then incorporating the micelle-forming copolymers in PBD- and PBA-based polymersomes the antiviral potency could be improved most likely due to the larger, virus-mimicking size and flexible membrane architecture of polymersomes. Multivalent interaction with SARS-CoV-2 spike proteins with HS-polymersomes was demonstrated on a single-molecule level using FCS, which is another factor contributing to their potency. Importantly, the HS-polymersomes revealed sub-nanomolar IC_50_’s against HS-dependent SARS-CoV-2 and HS-independent IAV (H1N1), while protecting against alphavirus (SINV) infection in vivo using the zebrafish embryo model. These findings position PBD-PM and PBA-PM as compelling candidates for further preclinical development and establish the HS-polymersome platform as a versatile entry inhibitor against emerging viral pathogens. Our potent virus-binding polymersomes open several opportunities for future development, including towards a virucidal system,^16,21–23^ co-delivery of antiviral drugs in the hydrophilic and hydrophobic compartments,^30,34^ and immune cell coordination.^31,32^

## Supporting Information

Supporting Information is available from:

## Acknowledgements

S.M. and A.N. thank the CDT of Translation Research in Advanced Cell and Gene Therapies at King’s College London for supporting this research. X.Z. and A.N. thank the Matt Wilson Scholarship from King’s College London for funding cryo-TEM imaging. I.W. thanks the Faculty of Engineering Widening Participation Fund and Department of Materials at Imperial College London for their support. A.N., R.P.G., and V.T. acknowledge support from King’ Together grants from King’s College London. A.N. and O.G-R. thank the Royal Society for their respective Research Grant (RGS\R2\242322 and RG\R1\251346). Z.P. and V.T. acknowledge support from the China Scholarship Council (CSC). V.T. is also supported by an MRC New Investigator Research Grant (MR/Z504178/1). Part of this research was funded by the UK Medical Research Council Discovery Awards to the G2P2 consortia MR/Y004205 to R.P.G. and W.S.B. We thank CUI at King’s College London and the Birkbeck, University of London (BBK) EM facility for cryo-TEM imaging. The TOC schematic and schematics in the Figures were generated by using and modifying Servier Medical Art (https://smart.servier.com/), provided by Servier, licensed under a Creative Commons Attribution 3.0 unported license.

## Author Contributions

S.M. performed chemical synthesis, particle formation and characterization, SARS-CoV-2 PV work, cellular toxicity work, and wrote the manuscript. R.Z., R.S., X.Z., F.K., I.W., and P.S.L.R. assisted particle formation and characterization. X.Z. and X.X. assisted SARS-CoV-2 PV assays. R.Z., R.S., J.C.B., K.Y.M, and R.K. performed the work with whole infectious SARS-CoV-2 and H1N1. Z.P. performed in vivo testing on SINV in the zebrafish embryo model. B.K. and O.R-G. synthesized and characterized PBA-PEG. V.T. supervised zebrafish embryo infection model. R.P.G supervised the SARS-CoV-2 PV work. W.B. supervised full virus work. A.N. designed, supervised and led the study and co-wrote the paper. All authors have read and commented on the manuscript.

## References

(1) Faramarzi, A.; Norouzi, S.; Dehdarirad, H.; Aghlmand, S.; Yusefzadeh, H.; Javan-Noughabi, J. The Global Economic Burden of COVID-19 Disease: A Comprehensive Systematic Review and Meta-Analysis. Syst. Rev. 2024, 13 (1), 68. 10.1186/s13643-024-02476-6

(2) World Health Organization (2023). *WHO Coronavirus (COVID-19) Dashboard*; 2025. https://data.who.int/dashboards/covid19/data

(3) Li, G.; Hilgenfeld, R.; Whitley, R.; De Clercq, E. Therapeutic Strategies for COVID-19: Progress and Lessons Learned. Nat. Rev. Drug Discov. 2023, 22 (6), 449–475. 10.1038/s41573-023-00672-y

(4) Symons, J.; Chung, C.; Verheijen, B. M.; Shemtov, S. J.; de Jong, D.; Amatngalim, G.; Nijhuis, M.; Vermulst, M.; Gout, J.-F. The Mutational Landscape of SARS-CoV-2 Provides New Insight into Viral Evolution and Fitness. Nat. Commun. 2025, 16 (1), 6425. 10.1038/s41467-025-61555-x

(5) World Health Organization. WHO Report Influenza (Seasonal). 2025.

(6) Zhao, T.; Chen, T.; Zhang, J.; Li, C.; Qin, C. Chikungunya Virus: Current Situation and Future Challenges. *Biosaf*. Health 2025, 7 (6), 348–356. 10.1016/j.bsheal.2025.10.001

(7) World Health Organization. Chikungunya Virus Disease-Global Situation. 2025.

(8) Karim, M.; Lo, C.-W.; Einav, S. Preparing for the next Viral Threat with Broad-Spectrum Antivirals. Journal of Clinical Investigation 2023, 133 (11). 10.1172/JCI170236

(9) Luong, Q. X. T.; Hoang, P. T.; Ho, P. T.; Ayun, R. Q.; Lee, T. K.; Lee, S. Potential Broad-Spectrum Antiviral Agents: A Key Arsenal Against Newly Emerging and Reemerging Respiratory RNA Viruses. Int. J. Mol. Sci. 2025, *26* (4), 1481. 10.3390/ijms26041481

(10) Pu, Y.; Zhu, C.; Liao, J.; Gong, L.; Wu, Y.; Liu, S.; Wang, H.; Zhang, Q.; Lin, Z. Antiviral Nanomedicine: Advantages, Mechanisms and Advanced Therapies. Bioact. Mater. 2025, 52, 92–122. 10.1016/j.bioactmat.2025.05.030

(11) Najer, A. Pathogen-binding Nanoparticles to Inhibit Host Cell Infection by Heparan Sulfate and Sialic Acid Dependent Viruses and Protozoan Parasites. Smart Medicine 2024, 3 (2). 10.1002/smmd.20230046

(12) Cagno, V. Challenges in Developing Broad-Spectrum Viral Attachment Inhibitors Targeting Heparan Sulfates and Sialic Acids. J. Virol. 2026, 100 (4). 10.1128/jvi.00139-26

(13) Cagno, V.; Tseligka, E. D.; Jones, S. T.; Tapparel, C. Heparan Sulfate Proteoglycans and Viral Attachment: True Receptors or Adaptation Bias? Viruses 2019, 11 (7), 596. 10.3390/v11070596

(14) Weis, W.; Brown, J. H.; Cusack, S.; Paulson, J. C.; Skehel, J. J.; Wiley, D. C. Structure of the Influenza Virus Haemagglutinin Complexed with Its Receptor, Sialic Acid. Nature 1988, 333 (6172), 426–431. 10.1038/333426a0

(15) Bhebhe, L. M.; Kim, J.; Jones, L. M.; Super, E. H.; Jones, S. T. Antiviral Mechanism Change of Poly(Styrene Sulfonate) through Gold Nanoparticle Coating. Polym. Chem. 2024, 15 (10), 945–951. 10.1039/D3PY01217D

(16) Cagno, V.; Andreozzi, P.; D’Alicarnasso, M.; Jacob Silva, P.; Mueller, M.; Galloux, M.; Le Goffic, R.; Jones, S. T.; Vallino, M.; Hodek, J.; Weber, J.; Sen, S.; Janeček, E.-R.; Bekdemir, A.; Sanavio, B.; Martinelli, C.; Donalisio, M.; Rameix Welti, M.-A.; Eleouet, J.-F.; Han, Y.; Kaiser, L.; Vukovic, L.; Tapparel, C.; Král, P.; Krol, S.; Lembo, D.; Stellacci, F. Broad-Spectrum Non-Toxic Antiviral Nanoparticles with a Virucidal Inhibition Mechanism. Nat. Mater. 2018, 17 (2), 195–203. 10.1038/nmat5053

(17) Groß, R.; Dias Loiola, L. M.; Issmail, L.; Uhlig, N.; Eberlein, V.; Conzelmann, C.; Olari, L.; Rauch, L.; Lawrenz, J.; Weil, T.; Müller, J. A.; Cardoso, M. B.; Gilg, A.; Larsson, O.; Höglund, U.; Pålsson, S. A.; Tvilum, A. S.; Løvschall, K. B.; Kristensen, M. M.; Spetz, A.; Hontonnou, F.; Galloux, M.; Grunwald, T.; Zelikin, A. N.; Münch, J. Macromolecular Viral Entry Inhibitors as Broad-Spectrum First-Line Antivirals with Activity against SARS-CoV-2. Advanced Science 2022, 9 (20). 10.1002/advs.202201378

(18) Najer, A.; Blight, J.; Ducker, C. B.; Gasbarri, M.; Brown, J. C.; Che, J.; Høgset, H.; Saunders, C.; Ojansivu, M.; Lu, Z.; Lin, Y.; Yeow, J.; Rifaie-Graham, O.; Potter, M.; Tonkin, R.; Penders, J.; Doutch, J. J.; Georgiadou, A.; Barriga, H. M. G.; Holme, M. N.; Cunnington, A. J.; Bugeon, L.; Dallman, M. J.; Barclay, W. S.; Stellacci, F.; Baum, J.; Stevens, M. M. Potent Virustatic Polymer–Lipid Nanomimics Block Viral Entry and Inhibit Malaria Parasites In Vivo. ACS Cent. Sci. 2022, 8 (9), 1238–1257. 10.1021/acscentsci.1c01368

(19) Nag, A.; Mukherjee, A.; Palit, S.; Jana, S. S.; Ghosh, S. Chain-Folding Regulated Assembly and Antiviral Activity of Sulfated Polyurethanes: A Structure–Property Relationship Study. ACS Appl. Polym. Mater. 2026, 8 (1), 555–563. 10.1021/acsapm.5c04030

(20) Nag, A.; Banerjee, K.; Barman, R.; Kar, J.; Sarkar, D. P.; Jana, S. S.; Ghosh, S. Direct Correlation between the Secondary Structure of an Amphiphilic Polymer and Its Prominent Antiviral Activity. J. Am. Chem. Soc. 2023, 145 (1), 579–584. 10.1021/jacs.2c11216

(21) Li, X.; Cheng, N.; Shi, D.; Li, Y.; Li, C.; Zhu, M.; Jin, Q.; Wu, Z.; Zhu, L.; He, Y.; Yao, H.; Ji, J. Sulfated Liposome-Based Artificial Cell Membrane Glycocalyx Nanodecoys for Coronavirus Inactivation by Membrane Fusion. Bioact. Mater. 2024, 33, 1–13. 10.1016/j.bioactmat.2023.10.021

(22) Li, Y.; Li, Y.; Li, X.; Liu, F.; Dai, W.; Chen, M.; Ji, J.; Jin, Q.; Wang, K. A Broad-Spectrum Antiviral Phase-Transformable Liposome with Membrane Fusion-Mediated Virucidal Capability. Adv. Healthc. Mater. 2025, 14 (20). 10.1002/adhm.202501446

(23) Martins, Y. A.; Bondeelle, L.; Zwygart, A. C.; de Lima, T. M.; Souza, J. de P.; Tee, H. K.; Chahud, F.; Neto, E. de A.; Stellacci, F.; Lopez, R. F. V.; Tapparel, C. Novel Wide-Spectrum Virucidal Lipid Nanoparticles. Small 2026, 22 (7). 10.1002/smll.202507669

(24) Mohammadifar, E.; Gasbarri, M.; Cagno, V.; Achazi, K.; Tapparel, C.; Haag, R.; Stellacci, F. Polyanionic Amphiphilic Dendritic Polyglycerols as Broad-Spectrum Viral Inhibitors with a Virucidal Mechanism. Biomacromolecules 2022, 23 (3), 983–991. 10.1021/acs.biomac.1c01376

(25) Dey, P.; Bergmann, T.; Cuellar-Camacho, J. L.; Ehrmann, S.; Chowdhury, M. S.; Zhang, M.; Dahmani, I.; Haag, R.; Azab, W. Multivalent Flexible Nanogels Exhibit Broad-Spectrum Antiviral Activity by Blocking Virus Entry. ACS Nano 2018, 12 (7), 6429–6442. 10.1021/acsnano.8b01616

(26) Khatri, V.; Boback, N.; Abdelwahab, H.; Niemeyer, D.; Palmer, T. M.; Sahoo, A. K.; Kerkhoff, Y.; Ludwig, K.; Heinze, J.; Balci, D.; Trimpert, J.; Haag, R.; Povolotsky, T. L.; Netz, R. R.; Drosten, C.; Lauster, D. C.; Bhatia, S. Polysialosides Outperform Sulfated Analogs for Binding with SARS-CoV-2. Small 2025, 21 (34). 10.1002/smll.202500719

(27) Etouki, R.; Xing, N.; Dimde, M.; Nie, C.; Donskyi, I. S.; Ludwig, K.; Mohammadifar, E.; Haag, R. Self-Assembled Nanosheets of Biocompatible Polymers as Universal Cell-Membrane Mimic to Block Viral Infection. Bioact. Mater. 2025, 52, 857–865. 10.1016/j.bioactmat.2025.06.035

(28) Phillips, B.; Trinh, T. K. H.; Beitzel, J.; Liu, S.; Hla, J.; Lin, X.; Chen, C.-L.; Zhang, M. Peptoid-Based Nanosheets Exhibiting Broad Antiviral Activity against Enveloped RNA Viruses. ACS Nano 2026, 20 (20), 14860–14878. 10.1021/acsnano.6c04885

(29) Cagno, V.; Gasbarri, M.; Medaglia, C.; Gomes, D.; Clement, S.; Stellacci, F.; Tapparel, C. Sulfonated Nanomaterials with Broad-Spectrum Antiviral Activity Extending beyond Heparan Sulfate-Dependent Viruses. Antimicrob. Agents Chemother. 2020, 64 (12). 10.1128/AAC.02001-20

(30) Nazemi, A.; Haeryfar, S. M. M.; Gillies, E. R. Multifunctional Dendritic Sialopolymersomes as Potential Antiviral Agents: Their Lectin Binding and Drug Release Properties. Langmuir 2013, 29 (21), 6420–6428. 10.1021/la400890f

(31) Zhu, Y.; Wei, L.; Zwygart, A. C.; Gaínza, P.; Khac, Q. O.; Olgiati, F.; Kurum, A.; Tang, L.; Correia, B.; Tapparel, C.; Stellacci, F. A Synthetic Multivalent Lipopeptide Derived from Pam3CSK4 with Irreversible Influenza Inhibition and Immuno-Stimulating Effects. Small 2024, 20 (31). 10.1002/smll.202307709

(32) Chen, J.; Zhang, G.; Liu, F.; Li, X.; Li, S.; Liu, Y.; Yang, C.; Pierleoni, G.; Li, H.; Li, Q.; Li, Z.; Tagliabue, A.; Chang, Y.-Z.; Wang, L.; Andreano, E.; Yu, P.; Li, Y. CuInP _2_ S _6_ Captures Virus for Viral Inhibition and Vaccine Adjuvanticity. Nano Lett. 2026, 26 (12), 4034–4045. 10.1021/acs.nanolett.5c04910

(33) Discher, B. M.; Won, Y.-Y.; Ege, D. S.; Lee, J. C.-M.; Bates, F. S.; Discher, D. E.; Hammer, D. A. Polymersomes: Tough Vesicles Made from Diblock Copolymers. Science (1979). 1999, 284 (5417), 1143–1146. 10.1126/science.284.5417.1143

(34) Chun, H.; Yeom, M.; Kim, H.-O.; Lim, J.-W.; Na, W.; Park, G.; Park, C.; Kang, A.; Yun, D.; Kim, J.; Song, D.; Haam, S. Efficient Antiviral Co-Delivery Using Polymersomes by Controlling the Surface Density of Cell-Targeting Groups for Influenza A Virus Treatment. Polym. Chem. 2018, 9 (16), 2116–2123. 10.1039/C8PY00116B

(35) Najer, A.; Kim, J.; Saunders, C.; Che, J.; Baum, J.; Stevens, M. M. Enhanced Antimalarial and Antisequestration Activity of Methoxybenzenesulfonate-Modified Biopolymers and Nanoparticles for Tackling Severe Malaria. ACS Infect. Dis. 2024, 10 (2), 732–745. 10.1021/acsinfecdis.3c00564

(36) Vonnemann, J.; Sieben, C.; Wolff, C.; Ludwig, K.; Böttcher, C.; Herrmann, A.; Haag, R. Virus Inhibition Induced by Polyvalent Nanoparticles of Different Sizes. Nanoscale 2014, 6 (4), 2353. 10.1039/c3nr04449a

(37) Farndale, R.; Buttle, D.; Barrett, A. Improved Quantitation and Discrimination of Sulphated Glycosaminoglycans by Use of Dimethylmethylene Blue. Biochimica et Biophysica Acta (BBA) - General Subjects 1986, 883 (2), 173–177. 10.1016/0304-4165(86)90306-5

(38) Barbosa, I. Improved and Simple Micro Assay for Sulfated Glycosaminoglycans Quantification in Biological Extracts and Its Use in Skin and Muscle Tissue Studies. Glycobiology 2003, 13 (9), 647–653. 10.1093/glycob/cwg082

(39) Najer, A.; Rifaie-Graham, O.; Yeow, J.; Adrianus, C.; Chami, M.; Stevens, M. M. Differences in Human Plasma Protein Interactions between Various Polymersomes and Stealth Liposomes as Observed by Fluorescence Correlation Spectroscopy. Macromol. Biosci. 2023, 23 (8). 10.1002/mabi.202200424

(40) Rifaie-Graham, O.; Yeow, J.; Najer, A.; Wang, R.; Sun, R.; Zhou, K.; Dell, T. N.; Adrianus, C.; Thanapongpibul, C.; Chami, M.; Mann, S.; de Alaniz, J. R.; Stevens, M. M. Photoswitchable Gating of Non-Equilibrium Enzymatic Feedback in Chemically Communicating Polymersome Nanoreactors. Nat. Chem. 2023, 15 (1), 110–118. 10.1038/s41557-022-01062-4

(41) de Kruijff, R. M.; Raavé, R.; Kip, A.; Molkenboer-Kuenen, J.; Roobol, S. J.; Essers, J.; Heskamp, S.; Denkova, A. G. Elucidating the Influence of Tumor Presence on the Polymersome Circulation Time in Mice. Pharmaceutics 2019, 11 (5), 241. 10.3390/pharmaceutics11050241

(42) Meeuwissen, S. A.; Bruekers, S. M. C.; Chen, Y.; Pochan, D. J.; van Hest, J. C. M. Spontaneous Shape Changes in Polymersomes via Polymer/Polymer Segregation. Polym. Chem. 2014, 5 (2), 489–501. 10.1039/C3PY00906H

(43) Paiardi, G.; Richter, S.; Oreste, P.; Urbinati, C.; Rusnati, M.; Wade, R. C. The Binding of Heparin to Spike Glycoprotein Inhibits SARS-CoV-2 Infection by Three Mechanisms. Journal of Biological Chemistry 2022, 298 (2), 101507. 10.1016/j.jbc.2021.101507

(44) Clausen, T. M.; Sandoval, D. R.; Spliid, C. B.; Pihl, J.; Perrett, H. R.; Painter, C. D.; Narayanan, A.; Majowicz, S. A.; Kwong, E. M.; McVicar, R. N.; Thacker, B. E.; Glass, C. A.; Yang, Z.; Torres, J. L.; Golden, G. J.; Bartels, P. L.; Porell, R. N.; Garretson, A. F.; Laubach, L.; Feldman, J.; Yin, X.; Pu, Y.; Hauser, B. M.; Caradonna, T. M.; Kellman, B. P.; Martino, C.; Gordts, P. L. S. M.; Chanda, S. K.; Schmidt, A. G.; Godula, K.; Leibel, S. L.; Jose, J.; Corbett, K. D.; Ward, A. B.; Carlin, A. F.; Esko, J. D. SARS-CoV-2 Infection Depends on Cellular Heparan Sulfate and ACE2. Cell 2020, 183 (4), 1043–1057.e15. 10.1016/j.cell.2020.09.033

(45) Najer, A.; Belessiotis-Richards, A.; Kim, H.; Saunders, C.; Fenaroli, F.; Adrianus, C.; Che, J.; Tonkin, R. L.; Høgset, H.; Lörcher, S.; Penna, M.; Higgins, S. G.; Meier, W.; Yarovsky, I.; Stevens, M. M. Block Length-Dependent Protein Fouling on Poly(2-oxazoline)-Based Polymersomes: Influence on Macrophage Association and Circulation Behavior. Small 2022, 18 (27). 10.1002/smll.202201993

(46) Rigler, P.; Meier, W. Encapsulation of Fluorescent Molecules by Functionalized Polymeric Nanocontainers: Investigation by Confocal Fluorescence Imaging and Fluorescence Correlation Spectroscopy. J. Am. Chem. Soc. 2006, 128 (1), 367–373. 10.1021/ja056719u

(47) Najer, A.; Wu, D.; Bieri, A.; Brand, F.; Palivan, C. G.; Beck, H.-P.; Meier, W. Nanomimics of Host Cell Membranes Block Invasion and Expose Invasive Malaria Parasites. ACS Nano 2014, 8 (12), 12560–12571. 10.1021/nn5054206

(48) Xiang, Q.; Li, L.; Wu, J.; Tian, M.; Fu, Y. Application of Pseudovirus System in the Development of Vaccine, Antiviral-Drugs, and Neutralizing Antibodies. Microbiol. Res. 2022, 258, 126993. 10.1016/j.micres.2022.126993

(49) Plante, J. A.; Liu, Y.; Liu, J.; Xia, H.; Johnson, B. A.; Lokugamage, K. G.; Zhang, X.; Muruato, A. E.; Zou, J.; Fontes-Garfias, C. R.; Mirchandani, D.; Scharton, D.; Bilello, J. P.; Ku, Z.; An, Z.; Kalveram, B.; Freiberg, A. N.; Menachery, V. D.; Xie, X.; Plante, K. S.; Weaver, S. C.; Shi, P.-Y. Spike Mutation D614G Alters SARS-CoV-2 Fitness. Nature 2021, 592 (7852), 116–121. 10.1038/s41586-020-2895-3

(50) Zhang, P.; Yan, H.; Liang, Z.; Zhang, P.; Li, X.-H.; Yuan, X.-Z.; Yu, G.; Wang, W.; Cai, C. Synthesis of Fucoidan-Biomimetic Glycopolymers with Flexible Skeletons for Enhanced Anti-Herpes Virus Efficacy. ACS Nano 2025, 19 (16), 15411–15424. 10.1021/acsnano.4c15060

(51) Dimova, R.; Seifert, U.; Pouligny, B.; Förster, S.; Döbereiner, H.-G. Hyperviscous Diblock Copolymer Vesicles. The European Physical Journal E 2002, 7 (3), 241–250. 10.1140/epje/i200101032

(52) Nie, C.; Sahoo, A. K.; Netz, R. R.; Herrmann, A.; Ballauff, M.; Haag, R. Charge Matters: Mutations in Omicron Variant Favor Binding to Cells. ChemBioChem 2022, 23 (6). 10.1002/cbic.202100681

(53) Kwon, P. S.; Xu, S.; Oh, H.; Kwon, S.-J.; Rodrigues, A. L.; Feroz, M.; Fraser, K.; He, P.; Zhang, F.; Hong, J. J.; Linhardt, R. J.; Dordick, J. S. Suramin Binds and Inhibits Infection of SARS-CoV-2 through Both Spike Protein-Heparan Sulfate and ACE2 Receptor Interactions. *Commun*. Biol. 2023, 6 (1), 387. 10.1038/s42003-023-04789-z

(54) Sullivan-Hill, B.; Shahrudin, S.; Ashley, C. L.; Shields, N. J.; Spijkers-Shaw, S.; Li, Y.; Devlin, R.; Rendle, P. M.; Wang, C.; Steain, M.; Zubkova, O. V.; Taylor, J. A.; Netzler, N. E. SARS-CoV-2 Variant-Specific Enhancement of Dendritic Heparan Sulfate Mimetic Antiviral Activities. Virology 2026, 620, 110915. 10.1016/j.virol.2026.110915

(55) Sofyantoro, F.; Sendi Priyono, D.; Indah Septriani, N.; Aristyaning Putri, W.; Mamada, S. S.; Adipuri Ramadaningrum, W.; Wijayanti, N.; Frediansyah, A.; Nainu, F. Zebrafish as a Model Organism for Virus Disease Research: Current Status and Future Directions. Heliyon 2024, 10 (13), e33865. 10.1016/j.heliyon.2024.e33865

(56) Passoni, G.; Langevin, C.; Palha, N.; Mounce, B. C.; Briolat, V.; Affaticati, P.; De Job, E.; Joly, J.-S.; Vignuzzi, M.; Saleh, M.-C.; Herbomel, P.; Boudinot, P.; Levraud, J.-P. Imaging of Viral Neuroinvasion in the Zebrafish Reveals That Sindbis and Chikungunya Viruses Favour Different Entry Routes. Dis. Model. Mech. 2017, 10 (7), 847–857. 10.1242/dmm.029231

(57) Boucontet, L.; Passoni, G.; Thiry, V.; Maggi, L.; Herbomel, P.; Levraud, J.-P.; Colucci-Guyon, E. A Model of Superinfection of Virus-Infected Zebrafish Larvae: Increased Susceptibility to Bacteria Associated With Neutrophil Death. Front. Immunol. 2018, 9 (MAY). 10.3389/fimmu.2018.01084

(58) Levraud, J.-P.; Palha, N.; Langevin, C.; Boudinot, P. Through the Looking Glass: Witnessing Host–Virus Interplay in Zebrafish. Trends Microbiol. 2014, 22 (9), 490– 497. 10.1016/j.tim.2014.04.014

(59) Mathez, G.; Silva, P. J.; Carlen, V.; Deloizy, C.; Prompt, C.; Hubart, A.; Rocha-Pereira, J.; Galloux, M.; Le Goffic, R.; Stellacci, F.; Cagno, V. A Pan-Respiratory Virus Attachment Inhibitor with High Potency in Human Airway Models and in Vivo. Sci. Adv. 2025, 11 (31), 9311. 10.1126/sciadv.adv9311

(60) Byrnes, A. P.; Griffin, D. E. Binding of Sindbis Virus to Cell Surface Heparan Sulfate. J. Virol. 1998, 72 (9), 7349–7356. 10.1128/JVI.72.9.7349-7356.1998

