## Supporting Information for "Antiviral Sulfonated Polymersomes Inhibit Corona-, Influenza-, and Alphavirus"

### Materials and Methods

#### HS-mimicking micelle and polymersome formulation

Poly(n-butyl acrylate)-*block*-poly(acrylic acid) (PBA-PAA) (7.5kDa-7.5kDa, Polymer Source, P42707A2-nBuAAA) and poly(DL-lactic acid)-*block*-poly(acrylic acid) (PDLLA-PAA) (9kDa-9kDa, Sigma-Aldrich, 802190) micelles were directly dispersed in MES buffer at a pH of 6 (0.5 M MES, 0.25 M NaCl, pH 6) at 10 mg/mL by stirring overnight. Poly(1,4 butadiene)-*block*-poly(acrylic acid) (PBD-PAA) (0.9kDa-2.6kDa, Polymer Source, P10508A-BdAA) and poly(dimethylsiloxane)-*block*-poly(acrylic acid) (PDMS-PAA) (8kDa-8kDa, Polymer Source, P43383A-DMSAA) were both dissolved in ethanol and assembled by the solvent injection method in a MES buffer at a pH of 6 (0.5 M MES, 0.25 M NaCl, pH 6) to yield 6.8 mg/mL and 10 mg/ml, respectively. Ethanol was evaporated off by blowing Argon gas on top of the stirred samples. Subsequent modifications followed a previous protocol.[1,2] The particles were modified to yield HS-micelles by adding 21.4 mg/mL 5-amino-2-methoxybenzene sulfonic acid (AMBS) (Chem Cruz, SC-233225) into each sample. This was followed by sequential addition of 6.6 mg 1-Ethyl-3-[3-dimethylaminopropyl] carbodiimide hydrochloride (EDC-HCl) (MilliporeSigma) in 12 µL MES pH 6 buffer per mL of sample every 30 - 60 min with 1 min sonication before each addition over a period of about 12 hours and a total of at least 10 additions. HS-micelles were then purified and buffer exchanged via sequential size exclusion chromatography (SEC). HS-micelles were first passed through a PD-MiniTrap G-25 column (Cytiva) equilibrated in a phosphate buffer (0.1 M phosphate, 0.05 M NaCl, pH 7.4) and further passed through PD-MidiTrap G-25 column (Cytiva) equilibrated in PBS. Once in PBS, HS-micelles were concentrated using Amicon Ultra 0.5mL Centrifugal Filters (MilliporeSigma) and passed through a 0.45 µm syringe filter to sterilize the particle suspensions before introduction into biological assays. Modification was confirmed based on Farndale *et al.* (Figure S4), using dimethylmethylene blue (DMMB) (15.8 µg/mL DMMB; 250

μL per well of a 96-well plate) against a poly(sodium 4-styrenesulfonate) (PSS, 1 MDa) standard (20-2.5 μg/mL).[3] 50 μL of PSS standard, modified nanoparticle sample in PBS, or PBS control was added to the DMMB. Absorbance spectra were recorded immediately after mixing, and the values at 590 nm were used to construct the PSS calibration curve and calculate the corresponding concentrations for the samples as we have done previously for similar nanoparticles.[1,2] Concentration values given in biological assays always represent the amount of active sulfonated polymer as obtained via DMMB assays and the corresponding nanoparticle concentrations as obtained by FCS as described below.

For HS-polymersome assembly, AMBS-modified copolymer micelles (PBD-PAA-AMBS and PBA-PAA-AMBS) after the EDC-HCl addition steps as described above were first passed through a PD-MidiTrap G-25 column equilibrated in a phosphate buffer (0.1 M phosphate, 0.05 M NaCl, pH 7.4) and subsequently a PD10 column run in $ddH_2O$. Cation exchange was performed via passing the particles in $ddH_2O$ through ca. 5 mL of packed BT AG 50W-XB resin ($H^+$ form, Bio-Rad) and eluted with the slow addition of 30 ml of $ddH_2O$. Samples were then concentrated in Amicon Ultra 0.5mL Centrifugal Filters (MilliporeSigma) and freeze dried. HS-polymersome samples were then formulated by mixing AMBS-modified copolymers PBD-PAA-AMBS and PBA-PAA-AMBS dissolved in ethanol with vesicle-forming copolymers poly(butadiene)-*block*-poly(ethylene glycol) (PBD-PEG) (2.5kDa-1.3kDa, P43224B-BdEO or 2.5kDa-1.5kDa, P19015-BdEO, Polymer Source) and poly(butyl acrylate)-*block*-poly(ethylene glycol) (PBA-PEG) (5.2kDa-2kDa),[4] respectively, following the film rehydration method. Typical amounts of copolymer used for each sample was 49 mg PBD-PEG with 4.9 mg PBD-PAA-AMBS and 20 mg PBA-PEG with 4.7 mg PBA-PAA-AMBS. Control PEG-only vesicles were made from the same amount of corresponding PEGylated copolymer only. Ethanol was evaporated off the single/blended copolymers via exogenous Argon gas and further desiccated for at least 2 hours to ensure complete evaporation of ethanol.

Film rehydration was then done with the addition of 600 μL phosphate buffer (0.1 M phosphate, 0.05 M NaCl, pH 7.4) and vigorous stirring for 16-24 hours or until all the copolymer has been dispersed. PBD-PM and PBD-PEG were extruded 15 times using an Avanti mini-extruder (MilllioporeSigma) through a 100 nm-pore membrane (Cytiva) to achieve a uniform distribution in particle size. HS-polymersomes were passed through a PD-MiniTrap G-25 column (Cytiva) equilibrated in phosphate buffer (0.1 M phosphate, 0.05 M NaCl, pH 7.4) and further passed through PD-MidiTrapTM G-25 column (Cytiva) equilibrated in PBS. HS-polymersomes were concentrated using Amicon Ultra 0.5 mL Centrifugal Filters (MilliporeSigma) to get particles to a working concentration and passed through a 0.45 μm filter to filter sterilize.

All particles were characterized via dynamic light scattering (DLS) with 70μL of sample in microcuvettes and samples were diluted 1:20 in 300 mM sucrose solution for zeta potential analysis using a Malvern Zetasizer. For cryo-TEM, 4 μL of sample was deposited on 200 mesh UltrAuFoil 1.2/1.3 (Quantifoil), excess liquid blotted away and vitrified into liquid ethane at -180 °C. Images were collected using a Falcon 3EC or Gatan K3 camera mounted on a Titan Krios G3i.

**FCS protein binding analysis**

Fluorescence correlation spectroscopy (FCS) was used to assess spike protein binding to the different polymersome formulations. Measurements were performed on mixtures of equal volumes of Omicron SARS-CoV-2 spike protein AF488 (AFG11109, R&D systems, spike-488) with 1:10 dilutions of polymersome stock samples in PBS. To obtain particle concentrations, samples were mixed with hydrophobic dye Bodipy630 before measurement. FCS measurements were conducted at 25 °C immediately after mixing using a commercial Zeiss LSM 780 or 880 confocal microscope (Carl Zeiss Microscopy, Jena, Germany) equipped with a 488 nm $Ar^+$ laser and a 633 nm HeNe laser for excitation. After calibration of the beam waist with a known fluorophore (OG488 or AF647), diffusion coefficients (D) could be calculated, which were then converted to hydrodynamic diameters through the Stokes-Einstein equation. A triplet fraction T with triplet time $\tau_T$ was applied for all fits, fixed between 1-10 µs. The structural parameter SP was always set to 5.

The data was analyzed using PyCorrFit version 1.1.7,[5] first with a one-component model to obtain polymersome concentrations (Bodipy630) and diffusion of unbound protein (AF488 or OG488) in the absence of polymersomes and then with a two-component fitting model for all the samples with protein. In the two-component model, the first component was fixed to freely diffusing labelled protein, with the diffusion time, $\tau_1$, obtained from the corresponding one-component fits. The second component represents the slow diffusing species with diffusion time, $\tau_2$. These fits yielded a % particle fraction and a corresponding diffusion time for this fraction, which was converted to hydrodynamic diameters as explained above. 2 out of 25 curves were excluded for the PBA-PM sample due to detecting aggregates (> 5 µm size).

The number of spike proteins per particle was calculated using the following formulas:

$$CPP = F_1 \times CPP_{protein} + (1 - F_1) \times CPP_{particle}$$

$$N_{protein\ per\ NP} = CPP_{particle}/CPP_{protein}$$

where $CPP$ is the measured counts per particle (in kHz), $CPP_{protein}$ was derived from the calibration measurements of free protein using one component fits and $F_1$corresponds to the free protein fraction from the two component fits. The equation was then solved for $CPP_{particle}$, which represents the signal per particle for the particle fraction corresponding to $(1 - F_1)$. $N_{protein\ per\ NP}$ is the number of proteins per particle presented in the figures. Control experiments for non-specific binding were performed analogously to the spike-488 experiments but using OG488-labelled human serum albumin (HSA-488) instead.[4]

**SARS-CoV-2 D614G pseudovirus production**

SARS-CoV-2 PV were produced via transfection of 2μg of HIV 8.91 Gag-Pol plasmid, 3μg of CSXW plasmid (HIV-firefly luciferase) and 2μg of D614G SARS-CoV-2 spike plasmid in the presence of 35 μg of PEI-Max (1 mg/ml) and OptiMEM (Gibco) in sub-confluent HEK293T cells transfected in 10 cm dishes and incubated at 37°C with a media change after 4 hours. Supernatant containing SARS-CoV-2 PV were harvested 72 hours post-transfection and were filtered through 0.45 μm filter.[6]

**SARS-CoV-2 D614G pseudovirus neutralization assay**

24 hours before inhibition assay start HeLa cell made to express entry receptor ACE2 (HeLa-ACE2, kindly provided by James Voss, Scripps Research, San Diego) were plated in 96-well tissue-culture plates at a cell density of 125,000 cells/mL to allow cells to adhere to the plate. Micelles and polymersomes were serially diluted to concentrations of interest and pre-incubated with PV at 37 °C before addition to the HeLa-ACE2 cells.[6] Cells incubated with micelles/polymersomes and PV were incubated for 48-hours before supernatant was removed and Steady-Glo reagent (Promega) was added to the cells. Luminescence was measured and values were normalized to infection control wells. GraphPad Prism (GraphPad Software, Version 11) was used to fit a non-linear regression curve to determine the $IC_{50}$ for each nanoparticle suspension.

**Whole infectious SARS-CoV-2 neutralization assay**

Whole infectious SARS-CoV-2 virus assays were conducted on Vero E6 cells (Nuvonis Technologies), which were cultured in DMEM (Gibco), supplemented with 10% (v/v) FBS, 1% (v/v) P/S and 1% (v/v) NEAA. Cells were seeded in 96-well plates at 20,000 cells per well and incubated overnight at 37 °C. In a separate 96-well plate, samples were two-fold or four-fold serially diluted in duplicates. Subsequently, 100 Tissue Culture Infectious Dose 50 ($TCID_{50}$) SARS-CoV-2 Omicron variant (BA.1 lineage, isolate M21021166) was added per well and incubated for 1 hour at 37 °C. Media was aspirated from the cell plate, and the virus/inhibitor mixtures were transferred onto the cells, followed by incubation at 37 °C for 48 hours. Cells were then fixed with 4% (v/v) paraformaldehyde (PFA, Gibco) in PBS for 30 minutes and treated with 0.6% (v/v) $H_2O_2$ in methanol for 20 minutes at RT. After washing with 0.05% (v/v) PBS-Tween, cells were incubated with rabbit mAb anti–SARS-CoV-2 nucleocapsid (Sino Biological, 40143-R019) 1:3000 in PBS 5% (w/v) skim milk powder for 1 hour at RT. Following four washes, sheep anti-rabbit HRP conjugate (MilliporeSigma, AP510P) 1:1000 in PBS 5% (w/v) skim milk and 1% (v/v) BSA was added for 1 hour. Plates were washed again after which TMB substrate (Europa Bioproducts) was added to plate for 20 minutes at RT in the dark and quenched with 0.5 M HCl. Absorbance was measured at 450 nm and 620 nm using a spectrophotometer (Vantastar Microplate Reader, BMG Labtech).

**Whole infectious IAV neutralization assay**

MDCK cells (5000 – 20,000 per well) were seeded in 96 well-plates and incubated overnight. Serial dilutions of PBD-PM and PBA-PM were prepared in 96-well dilution plates. Influenza A virus (IAV) (A/England/195/2009(H1N1); $100\times TCID_{50}$ in serum-free DMEM; 60 μL) was subsequently added to all wells. The virus and HS-polymersome mixtures were incubated at 37 °C for 1 hour. Culture medium was then aspirated from the seeded MDCK plate, and each corresponding virus and nanoparticle dilution was transferred onto the cells, and the plates were incubated at 37 °C for 18 hours.

Following infection overnight, the medium was removed and cells were fixed with of 4% (v/v) PFA for 30 min at room temperature. The fixative was removed and endogenous peroxidase activity was quenched with 0.6% (v/v) hydrogen peroxide in methanol for 20 min at room temperature. Plates were drained, blotted and washed twice with wash buffer (0.05% (v/v) Tween 20 in PBS). Mouse monoclonal anti-influenza A nucleoprotein antibody (Abcam, ab128193), diluted 1:3000 in 5% (w/v) skimmed milk powder in wash buffer, was added and incubated at 37 °C for 1 hour. Plates were washed four times with wash buffer, and horseradish peroxidase (HRP)-conjugated goat anti-mouse IgG secondary antibody (Abcam, ab205719), diluted 1:3000 in blocking buffer (5% (w/v) skimmed milk powder and 1% (w/v) BSA in wash buffer), was added and incubated at 37 °C for 1 hour. After four further washes, tetramethylbenzidine (TMB) substrate was added and the plates were developed in the dark at room temperature for 18 minutes. The reaction was stopped with 0.5 M HCl. Absorbance was read on a dual-wavelength program at 450 nm and 620 nm, and the 620 nm reference reading was subtracted from the 450 nm signal. Percentage infectivity was calculated relative to the PBS vehicle control, and dose–response curves were fitted to determine $IC_{50}$ values for each inhibitor.

**HS-micelle and polymersome cytocompatibility studies**

HS-micelle and polymersome toxicity was evaluated in HeLa-ACE2, Vero E6, and MDCK cells using the exact same setup, concentrations, and incubation times as for the viral assays above but without using any virus. Toxicity was measured using (3-(4,5-

dimethylthiazol-2-yl)-5-(3-carboxymethoxyphenyl)-2-(4-sulfophenyl)-2H-tetrazolium) (MTS) and phenazine methosulfate (PMS). After the incubation period, media was removed and MTS/PMS reagent diluted in phenol-red free RPMI was incubated with cells for 1-3 hours until a color change was visible and absorbance was read at 490 nm on a plate reader.

**SINV virus preparation**

Sindbis virus naked gRNA preparation: Construct T7-SINV-nsp3-mScarlet was kindly shared by the Castello lab. The plasmids were electroporated in *E. coli* DH5alpha, amplified and maxi-prepped according to standard procedures. 3 µg of plasmid was linearized with XhoI (1,5 µL, i.e., 30 units) in 1x CutSmart buffer and a total reaction volume of 30 µL. Reactions were incubated for 1h at 37 °C, and the enzyme was later inactivated by incubation at 80 °C for 30 min. 0.6 µg of linearized DNA from the reaction was then directly used as a template for in *vitro* transcription in a 20 µL reaction at 37°C, using HiScribe® T7 ARCA mRNA Kit (New England Biolabs), according to the manufacturer's guidelines, except for an extended incubation time of 1h. The DNA template was digested by treatment with 2 µL of DNase I solution (as provided by the kit) for 30 minutes. 1 µL of the solutions was loaded on a 1.5% Agarose gel (non-denaturing, ran at 100 V for approximately 20 min) containing SybrSafe (2 µL/50 µL agarose gel), and visualized via a transilluminator to ensure integrity. SINV RNA was then purified by Lithium chloride precipitation (provided by the HiScribe® T7 ARCA mRNA Kit). Briefly, 20 µL of the solution was treated with 25 µL of LiCl solution (as provided by the kit), incubated at -20 °C for 30 min and centrifuged at 4 °C for 15 min at max speed. Supernatant was pipetted out, and pellet was rinsed in 70% ethanol (500 µL). Samples were spun down at 4 °C for 10 min at max speed. Ethanol was then removed, pellets allowed to dry completely at RT (approx. 10 min) and finally resuspended in 20 µL of RNAse-free water. The pellet was encouraged to resolubilize by warming up at 60 °C in a heat block. Before being stored at -20°C. The protocol yielded mRNA at 120-130 ng/µL (20 µL). Sindbis virus preparation involved use of 1,000 ng (with PEI) of SINV RNA to transfect Vero E6 cells in duplicate 6-well plates. By 2 days post-transfection, cells developed visible cytopathic effect (CPE) and largely expressed mScarlet, as expected. Supernatant and infected cells were collected from infected cells at 3 days post-transfection and used to infect new Vero E6 cells plated in T75 flasks. The viral particles were capable of infecting new Vero E6 cells. The supernatant was collected from these cells at 4 days post-infection to prepare viral particles for the following experiments. The supernatant was filtered from cell debris, concentrated through a 20% sucrose cushion by ultracentrifugation, aliquoted, and frozen at -80°C.

**Zebrafish embryo infection model**

Zebrafish eggs (AB strain) were collected at 0 days post-fertilization (dpf), and sorted for viable fertilized eggs after extensive cleaning. Microbial load was reduced by bleaching the egg surface for 15 min (0.004% bleach in 0.5xE2 medium). Eggs were finally washed three times (sterile 0.5xE2 medium) and redistributed in 10 cm diameter Petri dishes (V = 25 mL/ dish) at 100 eggs/Petri dish. The developing embryos were kept in an incubator at 28.5°C. Dead or underdeveloped embryos were removed daily and fresh 0.5xE2 medium was added. Before commencing experiments (i.e. microinjections and microscopy observations), lack of movement was ensured. All experiments were terminated before 5 dpf. Zebrafish larvae are not protected under the Animal Scientific Procedure Act (ASPA) up to that stage.

SINV injections using T7-SINV-nsp3-mScarlet and handling of larvae were performed according to literature protocols,[7] with some modifications. At 2 dpf, dechorionated or spontaneously hatched larvae were anesthetized with 0.63 mM buffered tricaine (ethyl 3-

aminobenzoate methanesulfonate) in 0.5xE2 medium. The embryos were then aligned in a Petri dish coated with 1% agarose (in 0.5xE2 medium), soaked with anesthetic solution. Prior to injection, PBD-PM and PBA-PM stocks were mixed 3:1 with ~40 PFU/nL SINV-nsp3-mScarlet to yield a final concentration of ~10 PFU/nL SINV-nsp3-mScarlet and particle concentrations of 230 nM (950 µg/mL) and 170 nM (1750 µg/mL) for PBD-PM and PBA-PM, respectively. Larvae were then microinjected with 2 nL of this mixture using a capillary needle under a dissection microscope, yielding ~20 PFUs SINV-nsp3-mScarlet per larva. Injections were performed in the dorsal aorta. Larvae were recovered from anesthesia in 0.5xE2 medium, and groups of up to 20 larvae were distributed in individual wells of a 6-well plate, kept at 28.5 °C, and inspected daily up to 72 post-infection (hpi). For survival analysis, larvae failing to produce a heartbeat during a 30 s observation and that were unresponsive to touch, or movement of the plate were considered nonviable. Remains of dead embryos were removed from the wells. For live imaging, larvae were anaesthetized in buffered tricaine as above, positioned laterally on Petri dishes coated with 1% agarose (in 0.5xE2 medium), and covered with a film of anesthetic medium. Images of infected larvae were taken at 24 and 48 h hpi using a Leica MZ10 F fluorescence stereo microscope.

**Statistical analysis**

All the figure captions explain the sample sizes, employed statistical tests, and post hoc analysis. Each caption also clarifies data normalization and representation. All other data were analyzed and plotted using GraphPad Prism 11.0.0.

**Supporting Figures**

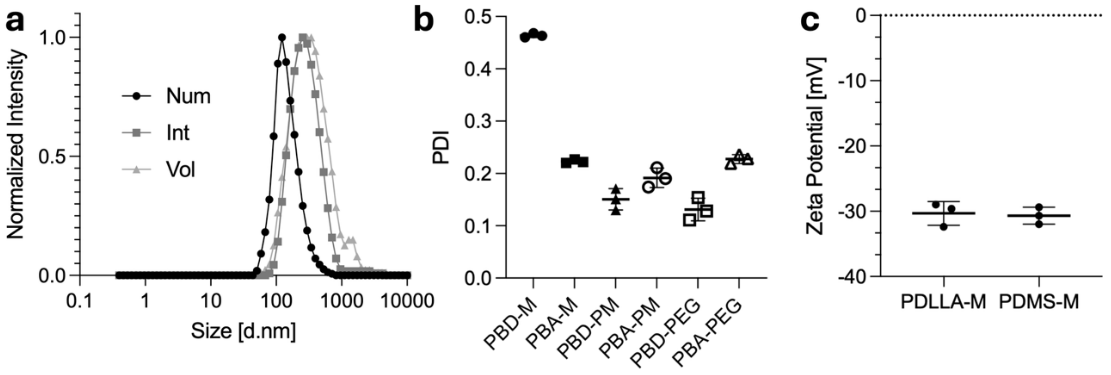


**Figure S1. Additional characterization of HS-micelles and HS-polymersomes.** (a) Representative DLS curves (intensity, number, and volume distribution) for PBA-PM (average of technical triplicates, number distribution from main Figure 1c is reproduced for comparison). (b) Polydispersity index (PDI) of each particle type derived from DLS, shown for micelles (PBD-M, PBA-M), polymersomes, (PBD-PM, PBA-PM), and the unmodified polymersomes (PBD-PEG, PBA-PEG) (mean ± SEM, technical triplicates). (c) Zeta potential average values (mean ± SEM, technical triplicates) for HS-micelles based on PDLLA and PDMS.

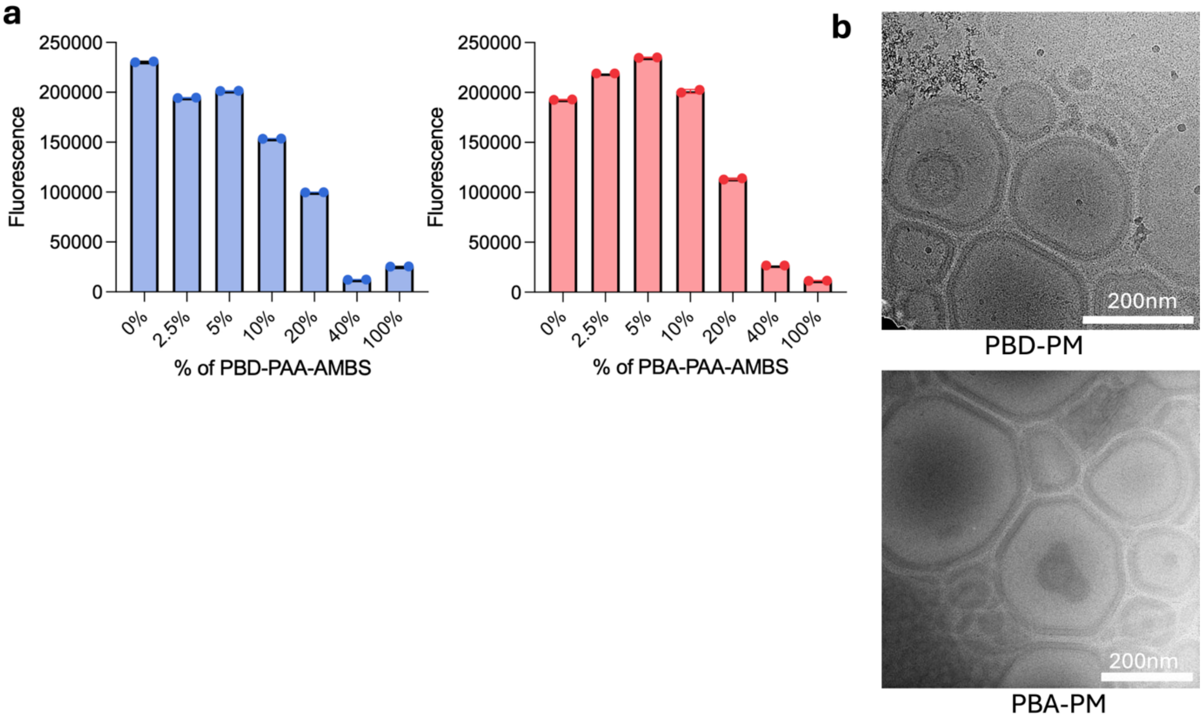


**Figure S2. SRB encapsulation and additional cryo-TEM images of polymersomes.** (a) Fluorescence after purification of SRB-loaded HS-polymersomes assembled at increasing ratios of modified (sulfonated) micelle-forming copolymer to vesicle-forming polymer (0, 2.5, 5, 10, 20, 40 and 100 mol%) for the PBD and PBA systems (technical duplicates). (b) Additional cryo-TEM images of the polymersomes PBD-PM and PBA-PM (scale bars 200 nm), confirming the vesicular morphologies.

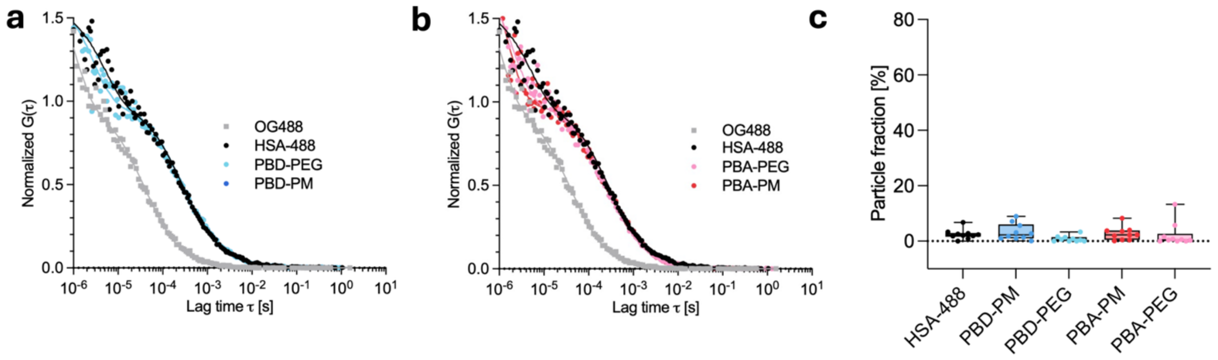


**Figure S3. Fluorescence correlation spectroscopy (FCS) of HSA binding to polymersomes.** (a, b) Average normalized autocorrelation curves for OG488-labeled HSA protein (HSA-488) and corresponding mixtures of HSA-488 with PBD-PEG, PBA-PEG, PBD-PM and PBA-PM ($n$ = 10 measurements). (c) Quantification of particle fraction for the slow diffusing species obtained from two-component fits of data in a,b ($n$ = 10 measurements).

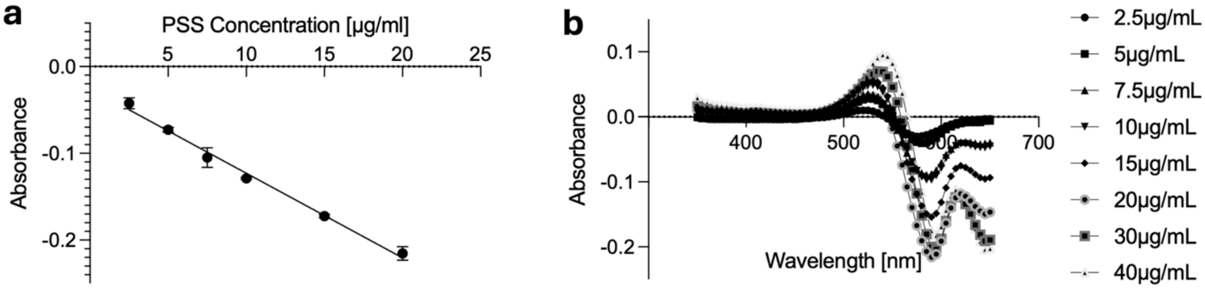


**Figure S4. Calibration curves of Farndale (DMMB) assay used to calculate nanoparticle sulfonation.** The sulfonated content of the HS-polymersomes and HS-micelles was assessed using the 1,9-dimethylmethylene blue (DMMB) assay, in which binding of the cationic dye to anionic sulfonate groups produces a concentration-dependent shift in absorbance at 590 nm, allowing the degree of sulfonation to be quantified against a sulfonated polymer standard.[3] (a) Standard curve of absorbance signal versus concentration of poly(styrene sulfonate) (PSS, 1 MDa) after subtraction of the DMMB/PBS blank (points represent mean ± SD of technical duplicates; line shows the linear regression). (b) Corresponding absorbance spectra across the PSS standard concentration series (2.5-40 µg/mL) after subtraction of the DMMB/PBS blank.

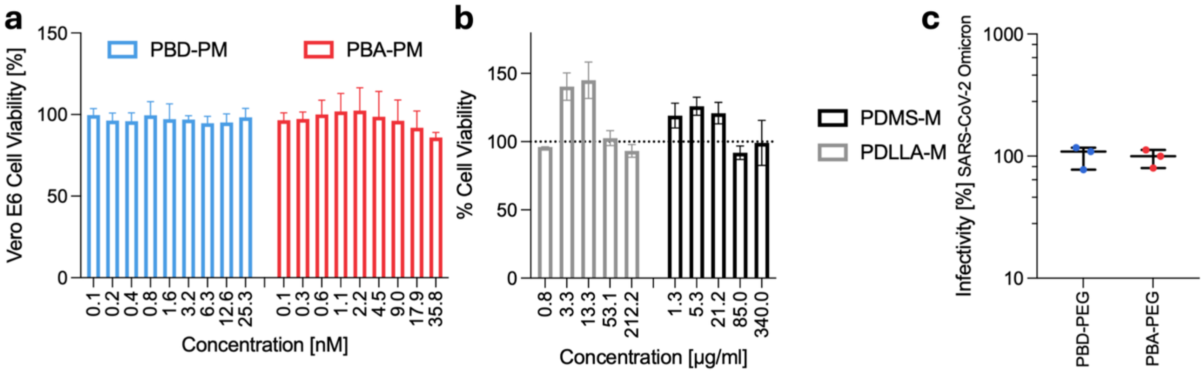


**Figure S5. Cytocompatibility and infectivity with nanoparticle treatment.** (a) Cell viability of Vero-E6 cells incubated with HS-mimetic nanoparticles for 48 h ($n \geq 3$ independent repeats with technical duplicates). (b) Cell viability of HeLa-ACE2 cells incubated with HS-mimetic nanoparticles for 48 h ($n$ = 1 independent repeat with technical duplicates). (c) Infectivity percentage for PBD-PEG and PBA-PEG vesicles alone for SARS-CoV-2 Omicron BA.1 matching the highest concentration of polymer used in main Figure 4 ($n$ = 3 independent repeats with technical duplicates).

### References


(1) Najer, A.; Kim, J.; Saunders, C.; Che, J.; Baum, J.; Stevens, M. M. Enhanced Antimalarial and Antisequestration Activity of Methoxybenzenesulfonate-Modified Biopolymers and Nanoparticles for Tackling Severe Malaria. *ACS Infect. Dis.* **2024**, *10* (2), 732–745. https://doi.org/10.1021/acsinfecdis.3c00564

(2) Najer, A.; Blight, J.; Ducker, C. B.; Gasbarri, M.; Brown, J. C.; Che, J.; Høgset, H.; Saunders, C.; Ojansivu, M.; Lu, Z.; Lin, Y.; Yeow, J.; Rifaie-Graham, O.; Potter, M.; Tonkin, R.; Penders, J.; Doutch, J. J.; Georgiadou, A.; Barriga, H. M. G.; Holme, M. N.; Cunnington, A. J.; Bugeon, L.; Dallman, M. J.; Barclay, W. S.; Stellacci, F.; Baum, J.; Stevens, M. M. Potent Virustatic Polymer-Lipid Nanomimics Block Viral Entry and Inhibit Malaria Parasites in Vivo. *ACS Cent. Sci.* **2022**, *8* (9), 1238–1257. https://doi.org/10.1021/acscentsci.1c01368

(3) Farndale, R. W.; Buttle, D. J.; Barrett, A. J. *Improved Quantitation and Discrimination of Sulphated Glycosaminoglycans by Use of Dimethylmethylene Blue*; 1986; Vol. 883.

(4) Najer, A.; Rifaie-Graham, O.; Yeow, J.; Adrianus, C.; Chami, M.; Stevens, M. M. Differences in Human Plasma Protein Interactions between Various Polymersomes and

Stealth Liposomes as Observed by Fluorescence Correlation Spectroscopy. *Macromol. Biosci.* **2023**, *23* (8). https://doi.org/10.1002/mabi.202200424

(5) Müller, P.; Schwille, P.; Weidemann, T. PyCorrFit-Generic Data Evaluation for Fluorescence Correlation Spectroscopy. *Bioinformatics* **2014**, *30* (17), 2532–2533. https://doi.org/10.1093/bioinformatics/btu328

(6) Pickering, S.; Wilson, H.; Bravo, E.; Perera, M. R.; Seow, J.; Graham, C.; Almeida, N.; Fotopoulos, L.; Williams, T.; Moitra, A.; Winstone, H.; Nissen, T. A. D.; Galão, R. P.; Snell, L. B.; Doores, K. J.; Malim, M. H.; Neil, S. J. D. Antibodies to the RBD of SARS-CoV-2 Spike Mediate Productive Infection of Primary Human Macrophages. *Nature Communications* **2024**, *15* (1). https://doi.org/10.1038/s41467-024-54458-w

(7) Levraud, J.-P.; Colucci-Guyon, E.; Redd, M. J.; Lutfalla, G.; Herbomel, P. In Vivo Analysis of Zebrafish Innate Immunity. In *Innate Immunity*; Humana Press: Totowa, NJ, 2008; pp 337–363. https://doi.org/10.1007/978-1-59745-570-1_20
